# Reconstructing heterogeneous metabolic trajectories of *E. coli* diauxie via a dynamical Maximum Entropy Principle

**DOI:** 10.64898/2026.05.28.728366

**Authors:** A. Ferrero-Fernández, F. Kratzl, L. Kelley, K.S. Korolev, D. Masoero, D. Segrè, I. Dukovski, D. De Martino

**Affiliations:** Biofisika Institute (CSIC, UPV/EHU), University of the Basque Country, 48940 Leioa, Spain; Bioinformatics Program, Faculty of Computing and Data Sciences, Boston University, Boston, Massachusetts 02215, USA; Department of Physics, Boston University, Boston, Massachusetts 02215, USA; Biological Design Center, Boston University, Boston, Massachusetts 02215, USA; Grupo de Física Matemática, Instituto Superior Técnico, Universidade de Lisboa, Av. Rovisco Pais, 1049-001, Lisboa, Portugal; Centro de Estudos Florestais, Instituto Superior de Agronomia, Universidade de Lisboa, Tapada da Ajuda, 1349-017, Lisboa, Portugal; Department of Biology, Boston University, Boston, Massachusetts 02215, USA; Department of Biomedical Engineering, Boston University, Boston, Massachusetts 02215, USA; Center for Advanced Interdisciplinary Research, Ss. Cyril and Methodius University, Skopje, N. Macedonia; Ikerbasque and Fundacion Biofisika Bizkaia (CSIC, UPV/EHU), 48940 Leioa, Spain

## Abstract

The glucose–acetate diauxic shift in E. coli is classically described as an abrupt, population-wide switch from glucose to acetate consumption. Recent experiments challenge this view, revealing a robust intermediate regime of co-consumption whose single-cell basis remains unresolved: does it reflect coexisting specialized subpopulations, or genuine mixed metabolic states within individual cells? We first develop a two-state consumer–resource model in which cells optimally grow on either glucose or acetate, and show that observed co-consumption trajectories cannot be decomposed into convex combinations of the two subpopulations — ruling out discrete metabolic states as a sufficient explanation. Physico-chemical constraints of the metabolic network instead enforce genuine single-cell co-consumption across a continuous spectrum of phenotypes. To resolve this, we apply a dynamical maximum entropy (maximum caliber) framework constrained by batch and chemostat experiments, inferring time-resolved distributions of metabolic fluxes that naturally predict a continuum of single-cell phenotypes spanning glucose overflow, mixed substrate utilization, and acetate consumption — revealing co-consumption as a dominant, persistent feature of single-cell metabolism around the switch rather than an artifact of population averaging. Finally, we formulate a continuous consumer–resource model over metabolic state space, in which selection, phenotypic diffusion, and moving metabolic boundaries driven by environmental feedback reproduce single-cell co-consumption trajectories and complex dynamical trends inaccessible to discrete models. Together, our results recast diauxic adaptation as a continuous redistribution of single-cell metabolic states rather than a discrete switch.

## INTRODUCTION

Growth of *Escherichia coli* on glucose is among the most deeply explored phenomena in biology [1], having guided the discovery of fundamental principles of genetic and metabolic regulation—from the lac operon to catabolite repression [2, 3]. Over the decades, metabolism has come to exemplify how cells allocate limited resources, balance growth and efficiency, and adapt to changing environments [4].

Despite this long history, key aspects of metabolic adaptation remain unresolved even in this canonical setting. When multiple carbon sources are available, cells must decide which substrates to consume, how to allocate enzymatic and proteomic resources, and when to reorganize their metabolic network; yet microbes are also known to co-consume substrates under a variety of conditions, blurring the classical picture of sequential substrate preference [5]. The glucose–acetate diauxic shift in *E. coli* epitomizes this complexity [6] and is peculiar in that the “second” substrate, acetate, is not only externally supplied but also endogenously produced via overflow metabolism during growth on glucose. Overflow metabolism leads to acetate secretion as a consequence of proteome allocation and respiratory constraints [7, 8], so that cells effectively experience a dynamically evolving mixture of glucose and acetate even when initially supplied with glucose alone. Upon glucose depletion, cells transition to acetate consumption, a process shaped by thermodynamic control of the Pta–AckA pathway, oxygen availability, and global regulation [9, 10]. While this transition has traditionally been depicted as an abrupt, population-wide metabolic switch—with all cells uniformly entering a lag phase before synchronously switching to acetate—its mechanistic underpinnings remain only partially understood.

A growing body of evidence indicates that adaptation to new metabolic conditions is neither instantaneous nor uniform across cells. Biological processes are intrinsically stochastic, and although regulatory networks buffer much of this variability, fluctuations can be amplified to generate phenotypic heterogeneity within clonal populations. Such heterogeneity is increasingly recognized as a functional feature of microbial physiology, contributing to robustness, adaptability, and survival in fluctuating environments [11–13]. In the context of diauxic growth, single-cell experiments have revealed sustained growth, asynchronous responses, and long-lived variability during nutrient shifts [14–16].

Recent experimental and theoretical work has challenged the textbook picture of diauxie as population-wide, sequential substrate utilization. Several studies have reported a robust phase of glucose and acetate co-consumption during growth [9, 10, 17], raising a fundamental question: does co-consumption reflect a mixture of specialized subpopulations—some growing on glucose, others on acetate—as suggested by discrete population models [18], or does it instead arise from intermediate metabolic states within individual cells? Distinguishing between these scenarios is essential for understanding how metabolic networks constrain and shape phenotypic diversity.

Addressing this question requires frameworks that accommodate both network-level constraints and phenotypic variability. Consumer–resource and kinetic models have provided valuable insight into population dynamics and substrate utilization, including early dynamic flux balance approaches to diauxic growth [19, 20]. However, these models typically assume a small number of predefined metabolic states and offer limited access to the internal organization of metabolism. Constraint-based approaches such as Flux Balance Analysis (FBA) exploit genome-scale metabolic reconstructions [21] to predict feasible flux distributions under physicochemical constraints [22]. Extensions such as Flux Variability Analysis [23] have shown that many alternative flux configurations can support near-optimal growth, highlighting an intrinsic degeneracy of metabolic states relevant to analyzing system responses to perturbations [24].

The maximum entropy (MaxEnt) formulation of constraint-based metabolism builds directly on this observation [25, 26]. Rather than selecting a single optimal flux state, MaxEnt infers a probability distribution over all feasible metabolic states consistent with experimental constraints, providing a principled way to describe metabolic variability without introducing *ad hoc* noise [27]. This framework has been applied to infer flux distributions in steady-state and continuous cultures and to analyze population heterogeneity [28–30], and several variants have been developed to model dynamics [31, 32] and predict fluxes [33, 34]. When extended to dynamics through the principle of maximum caliber [31, 35], it offers a natural language for describing stochastic trajectories and time-dependent metabolic adaptation.

Here we integrate multiple modeling frameworks and experimental data to dissect the origins of metabolic heterogeneity during the *E. coli* diauxic shift. We begin by fitting a two-state consumer–resource model—in which cells either consume glucose or acetate—to batch and chemostat data, and develop a constraint-based decomposition test showing that observed co-consumption trajectories lie outside the convex hull of two optimally growing subpopulations. We then introduce a dynamical maximum entropy (maximum caliber) framework that infers time-dependent flux distributions from OD growth curves, naturally predicting a continuum of single-cell phenotypes and a rich exometabolome during the diauxic transition. Extending this to single-cell resolution by combining flow cytometry with constraint-based inference, we reconstruct intracellular flux distributions conditioned on individual growth rates and identify coexisting co-consuming metabolic phenotypes. Finally, we reconcile the statistical and dynamical perspectives through a continuous consumer–resource model over metabolic state space, in which phenotypic diffusion and moving boundary conditions—induced by changing nutrient concentrations—generate the observed population dynamics, including single-cell co-consumption trajectories and lineage-resolved metabolic histories.

## RESULTS

### Two-state discrete consumer–resource model

Our aim is to model quantitatively the acetate switch of *E. coli* —the transition from glucose consumption with acetate overflow to acetate consumption upon glucose depletion—with particular focus on the co-consumption regime. We start from the simplest non-trivial setting able to capture co-consumption: a two-subpopulation consumer–resource model in the spirit of [18]. The motivation is twofold. First, this discrete picture is the natural baseline against which heterogeneity-based explanations should be tested, since it embodies the standard interpretation of co-consumption as a population-level mixture of specialized cells. Second, by fitting the model to our batch and chemostat data and asking whether observed co-consumption trajectories can be decomposed into contributions from two optimally growing subpopulations, we can sharply identify the regime in which the discrete picture breaks down and a continuous singlecell description is required—thereby motivating the maximum entropy and continuous consumer–resource frameworks developed in the subsequent sections.

The culture is modeled as a mixture of two subpopulations: one taking up glucose and secreting acetate via overflow, the other consuming acetate. Consumer–resource models are a natural default here because they provide a minimal yet mechanistic description of how populations and substrates jointly evolve via mass balance, are straightforward to fit to standard growth measurements, and allow transparent coupling with metabolic-network–derived yields [18, 19]. The model, formalized as a dynamical system tracking subpopulation densities and substrate concentrations, is described in Appendix A and includes a threshold-based acetate overflow mechanism and transition rates between the two subpopulations.

As in [18], biomass yields are set by flux balance analysis on the genome-scale metabolic network model iAF1260 [21]. These yields are computed once at the outset—separately for each subpopulation under its single-substrate FBA optimum—and used as fixed parameters, rather than being recomputed at each time step as in a fully fledged dynamical FBA (dFBA) approach [19]. This assumption implies that each subpopulation maximizes its growth rate given its substrate.

Model parameters were fitted to a dataset spanning multiple values of the feed concentrations *G*_0_, *A*_0_, and the dilution rate *D*, encompassing both batch (*D* = 0) and chemostat (*D* ≠ 0) conditions, using statistical inference with experimentally informed priors (see Appendix A).

The model is sketched in Fig. 1A. Inferred uptake and overflow parameters are consistent with previous results [8, 9, 18]. The fitted transition rates are strongly asymmetric: switching from glucose to acetate feeding is fast (τ_*g→a*_ ∼ 1 min) while the reverse is slow (τ_*a→g*_ ∼ 1 day), triggered when the acetate-to-glucose concentration ratio reaches ∼ 10^2^ . In Fig. 1B we report a typical batch experiment and its model fit, finding good agreement. The inferred model makes semiquantitative predictions on independent pulse and step-change experiments (Appendix A), and linear stability analysis provides analytical insight into the dynamical response (Appendix A). However, as shown in Fig. 1C, even at a low dilution rate (*D* = 0.1 h^−1^ ) the model fails to capture the approach to steady state quantitatively: the experimental relaxation displays a pronounced overshoot in bacterial density followed by a slow decay, whereas the model predicts a smoother, monotonic trajectory. Higher dilution rates produce even more complex behavior. As shown below, these discrepancies are naturally resolved by the continuous consumer–resource framework (Fig. 4D).

**FIG. 1.**
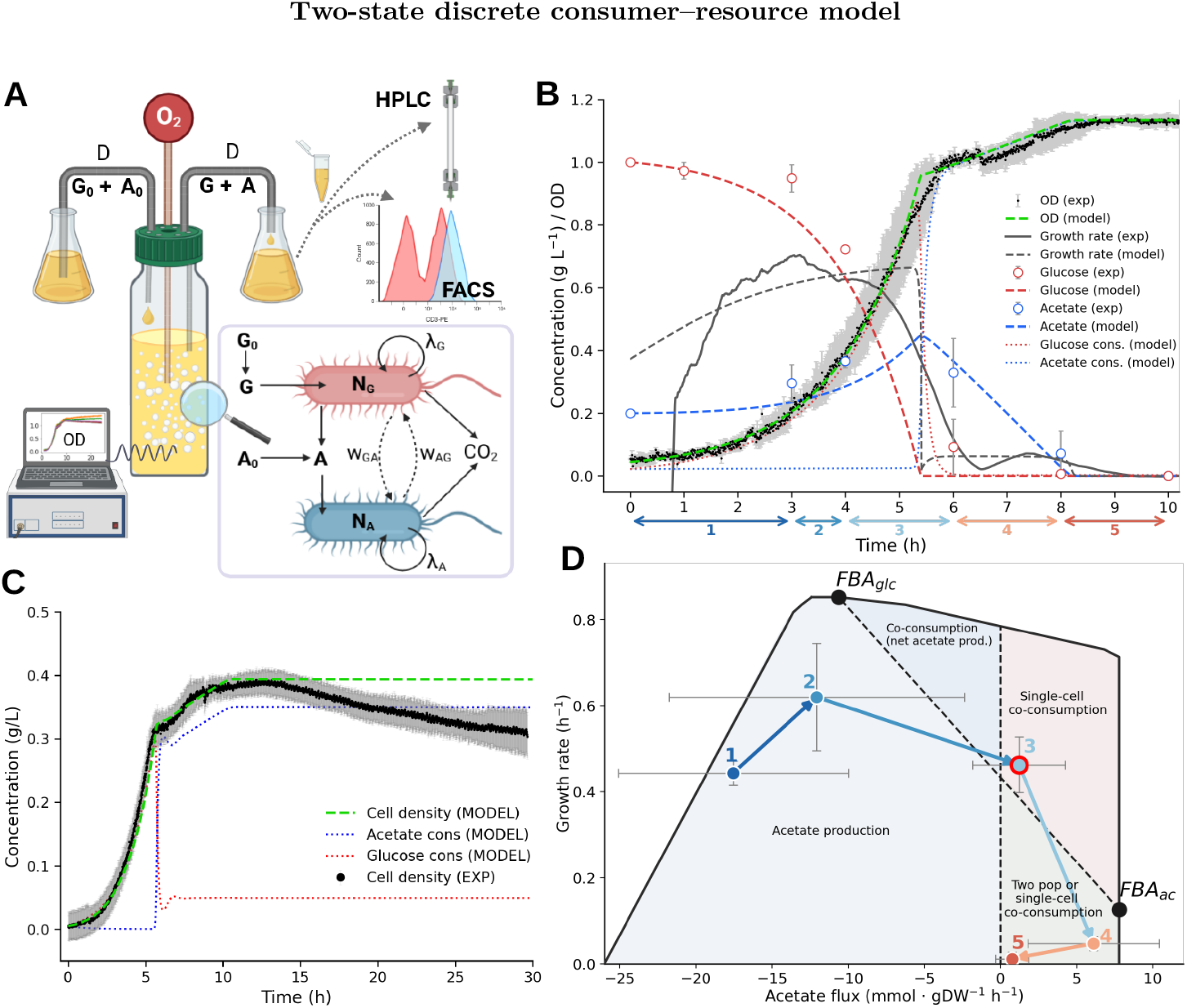
Optimal two-state consumer–resource model of the diauxic shift. **A**. Sketch of the experimental setting and of the model. **B**. Simulated and experimental curves used for the fitting; batch culture. Concentrations are in g L^−1^ and the growth rate λ is in h−1 . **C**. Model prediction of the OD curves under chemostat conditions of *D* = 0.1 h^−1^ . **D**. Projection of experimental growth rate and acetate flux values onto the feasible metabolic space from constraint-based network modeling. Negative and positive fluxes indicate secretion and uptake, respectively. Points FBA_ac_ and FBA_glc_ are the states of maximal growth for acetate and glucose feeders. Points above the segment connecting these two points cannot be decomposed into contributions from two separate subpopulations exchanging acetate.

Beyond these dynamical failures, the two-state picture is also problematic around the switch itself, where co-consumption is most prominent. To investigate this quantitatively, we developed a constraint-based decomposition test. Using a genome-scale metabolic network model, we computed via FBA the maximum-growth states **v**_*g*_ and **v**_*a*_ for glucose-only and acetate-only feeders, respectively. Within the two-state model, all feasible population states correspond to convex combinations on the segment **v**_*t*_ = *t* **v**_*g*_ + (1 − *t*) **v**_*a*_, *t* ∈ [0, 1]. We then swept the acetate flux across its full range and performed growth-rate maximization at each fixed acetate flux, tracing the boundary of the projected flux polytope in the (growth rate, acetate flux) plane (Fig. 1D). The FBA_glc_ vertex lies on this boundary at the maximum growth rate; FBA_ac_ lies on the positive acetate axis but inside the boundary, since a higher growth rate is achievable if some glucose uptake is permitted.

By construction, experimental points above the segment **v**_*g*_–**v**_*a*_ cannot be decomposed into two subpopulations exchanging a single substrate. Each point on the segment represents a convex combination of the two growth-rate-maximizing flux vectors, and therefore attains the highest growth rate that any such mixture of the two specialist subpopulations can achieve. Therefore, points above the segment represent states of single-cell co-consumption. From the experimental data shown in Fig. 1B, we computed five data points corresponding to successive measurement intervals around the switch, shown on the time axis in Fig. 1B (Appendix A). We found that the co-consumption trend (point 3 in Fig. 1D and time interval 3 in Fig. 1B), falls squarely in this undecomposable region. To investigate the possibility of presence of intermediate metabolic states within individual cells in the co-consumption time interval, in the following sections we will relax the growth rate optimality hypothesis, allowing cells to occupy mixed, suboptimal metabolic states via a dynamical extension of the maximum entropy method.

### Dynamical maximum entropy

In the previous section we have found that experimental data on the regime of co-consumption cannot be quantitatively reproduced by assuming two distinct subpopulations optimally feeding on glucose and exchanging acetate. One could extend the consumer–resource framework to three or more discrete states—for instance, adding an explicit co-consuming subpopulation—but this reintroduces the same conceptual problem at a finer level: any finite set of predefined, optimally growing states imposes an arbitrary discretization of what is an intrinsically high-dimensional continuum of feasible metabolic phenotypes. We therefore take the opposite route and let the data select which metabolic states are populated, allowing cells to occupy intermediate mixed regimes in which both substrates are consumed simultaneously.

To test this hypothesis, we apply the maximum entropy principle, generalized by Jaynes from statistical physics [36] and now a cornerstone of statistical inference widely used in biological data analysis [26, 37–41]. In the context of constraint-based metabolic modeling, maximum entropy has been used to model flux distributions in the presence of single-cell growth rate fluctuations [25, 26, 28] and extended to single-cell heterogeneity settings from gene expression [34, 42] to exchange interactions [43–45]. The approach imposes the underlying physicochemical constraints together with the requirement that the average growth rate matches the experimental value. This single constraint, combined with maximum entropy, dictates a Boltzmann–Gibbs flux distribution

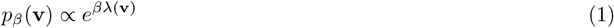

where *β* is set to match the experimental mean growth rate, ⟨λ⟩ = λ_exp_, and **v** is the metabolic state represented as a vector of reaction fluxes. Sampling from this distribution is more demanding than linear FBA optimization; however, sampling high-dimensional convex polytopes is nowadays well understood, with fast heuristics and robust software available [46–53]. Our approach is illustrated in Fig. 2A. We extended the static framework to dynamical settings by constructing a time series of Lagrange multipliers *β*(*t*) that solve the time sequence of variational problems

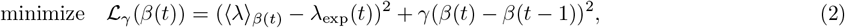

where γ is a regularizer set by cross-validation on a held-out test set (Appendix C). With it, we secure not only the match between the experimental and model average growth rates, but we also minimize the deviation of the Lagrange multipliers in consecutive time steps. At each time step, the sampled averages of growth rate and uptakes update the chemostat ODEs for external metabolite concentrations *c*_*i*_ and bacterial density *ρ*,

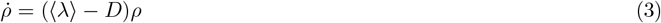

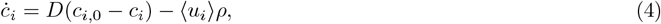

where *D* is the chemostat dilution rate, and ⟨*u*_*i*_⟩ are the average uptakes, bounded by Michaelis–Menten functions of concentrations (Appendix C), thus closing the dynamical inference loop.

**FIG. 2.**
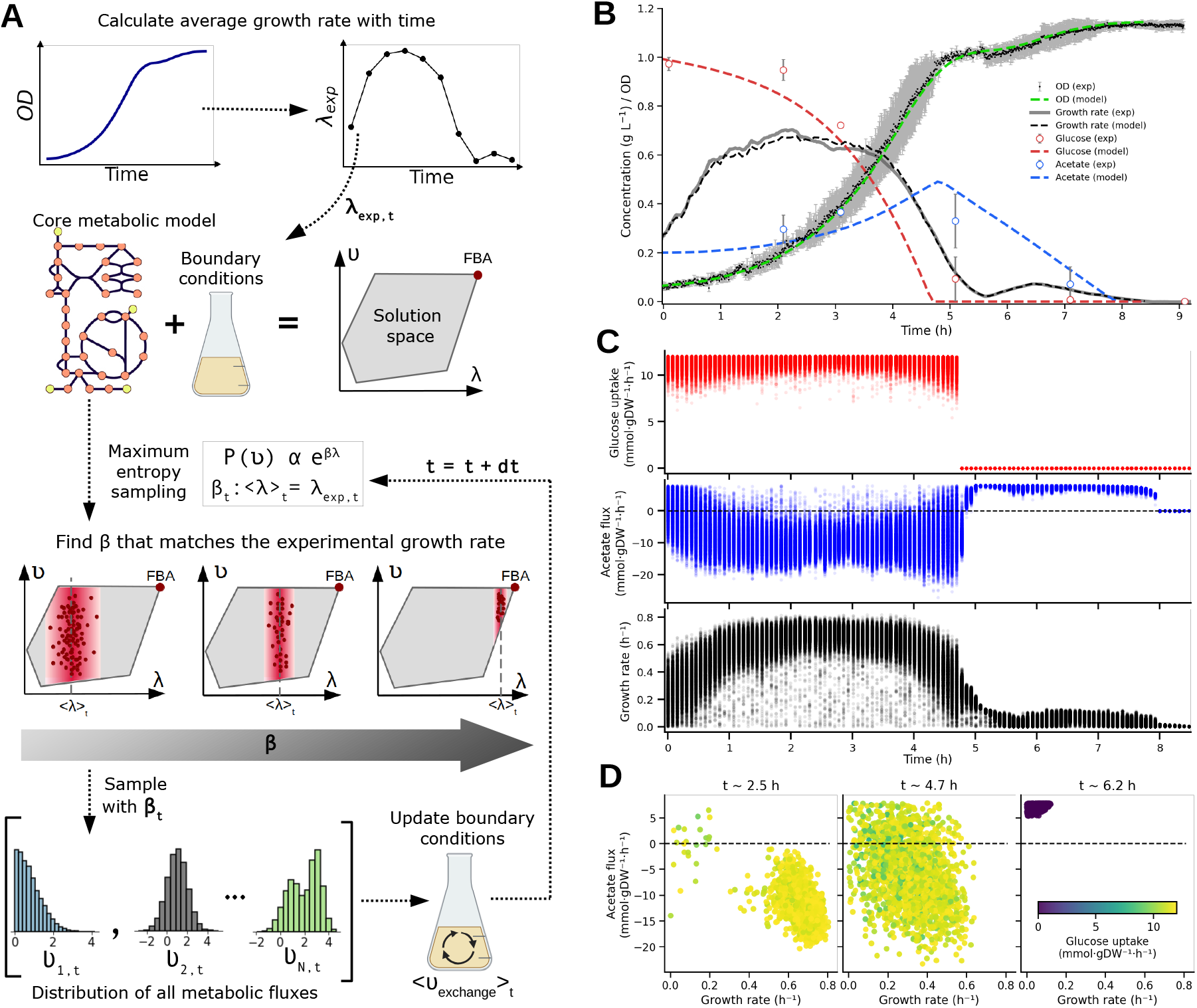
Dynamical maximum entropy approach. **A**. Schematic of the dynamical maximum entropy (maximum caliber) inference algorithm, applied to the same experimental setting depicted in Fig. 1A. From top to bottom: the time-dependent average growth rate is estimated from the derivative of the OD curves. At each time point, a core metabolic model of E. coli is constrained using the corresponding medium composition. The maximum entropy principle is then applied to sample the feasible flux space while fitting the observed average growth rate by tuning the parameter *β*. The resulting flux distributions are used to compute average exchange fluxes, which update the medium composition for the next time step. **B**. Comparison between experimental data and model predictions obtained using the approach in **A**. Trajectories are shown for metabolite concentrations, OD, and growth rate. **C**. Time-resolved distributions of selected fluxes from the simulation in **B**. From top to bottom: glucose uptake flux (mmol·gDW^−1^·h^−1^), acetate exchange flux (mmol·gDW^−1^·h^−1^; positive for uptake, negative for secretion), and growth rate (h^−1^). **D**. Scatter plots showing the relationship between growth rate, glucose uptake, and acetate exchange at three representative time points corresponding to before, during, and after the diauxic shift.

Model predictions are compared to experimental measurements in Fig. 2B. The method accurately captures the temporal trajectories of bacterial density and growth rate across the entire experiment, including the diauxic transition, and predicting trends of extracellular metabolite concentrations. Beyond acetate, the model predicts a broad exometabolome (see the appendix) comprising lactate, ethanol, succinate, and several organic acid intermediates— potentially explaining the minor peaks commonly observed in mass spectrometry data [54]. At the intracellular level, the model predicts continuous phenotypic distributions for all enzymatic fluxes in the catabolic carbon core, with acetate uptake and secretion exhibiting wide distributions and non-trivial correlations with glucose uptake and growth rate (Fig. 2C).

The time-resolved flux distributions reveal that around the metabolic switch most cells are predicted to operate in a regime of single-cell co-consumption of glucose and acetate, while exclusive consumption of a single substrate is largely confined to the earliest and latest hours of the experiment. These correlations are further illustrated in Fig. 2D, which shows scatter plots of growth rate, glucose uptake, and acetate exchange at representative time points before, during, and after the diauxic shift. The improved agreement with the data relative to the two-state discrete model is not merely a consequence of added fluctuations around two optimal states, but reflects the presence of genuinely co-consuming single-cell phenotypes lying in the undecomposable region identified in Fig. 1D—consistent with the decomposition test of the previous section. Having demonstrated that the maximum entropy method naturally populates the full metabolic space compatible with the data, we now extend it to single-cell resolution.

### Single-cell metabolic flux inference

The framework of the previous section suggests that cells occupy a continuum of metabolic flux phenotypes. To test this at single-cell resolution, we combine population-level experimental measurements of metabolite concentrations and bacterial density with single-cell flow cytometry data and maximum entropy constraint-based modeling [32, 34, 42]. From flow cytometry we extract the growth rate of cell *i* by assuming exponential growth,

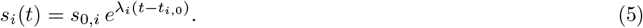

The scale is set by matching the population-average growth rate to the average log cytometry intensity; uncertainties are estimated by propagating the maximal errors in initial cell size and cell-cycle phase. Individual growth rates are then included as constraints in the metabolic network model, while the population-average glucose and acetate uptakes are matched by Boltzmann distributions with Lagrange multipliers *β*_*g*_, *β*_*a*_. The problem is to sample, for each cell *i*, a flux vector 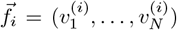 (the vector of all *N* intracellular reaction rates) from the joint distribution over all *N*_*c*_ ∼ 10^4^ cells,

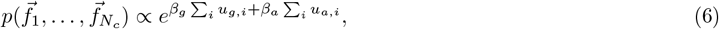

where *u*_*g,i*_ and *u*_*a,i*_ denote the glucose and acetate exchange fluxes of cell *i* (components of 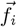), subject to the constraints that each flux vector lies in the stoichiometric flux polytope *P* (defined by 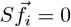 and global flux bounds) and that the growth rate of each cell lies within a narrow window around its experimentally inferred value,

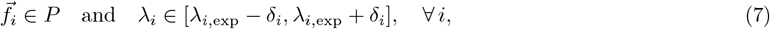

where λ_*i*,exp_ is the growth rate of cell *i* inferred from its FSC signal, and *δ*_*i*_ is the associated uncertainty propagated from the cell-size measurement. The Lagrange multipliers *β*_*g*_, *β*_*a*_ are then determined by requiring that the population-averaged exchange fluxes match the experimentally measured bulk uptake rates,

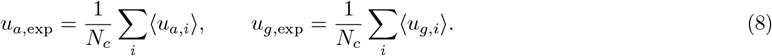

Figure 3 summarizes the single-cell metabolic flux inference. Flow cytometry measurements at selected time points quantify single-cell size distributions (Fig. 3A); applying the exponential growth law converts these into growth-rate distributions at the single-cell level [55]. These inferred growth rates are fixed as constraints for each cell, and maximum entropy inference is applied to sample the feasible intracellular flux space while matching the population-average glucose and acetate uptake rates via two Lagrange multipliers, *β*_glc_ and *β*_ac_.

**FIG. 3.**
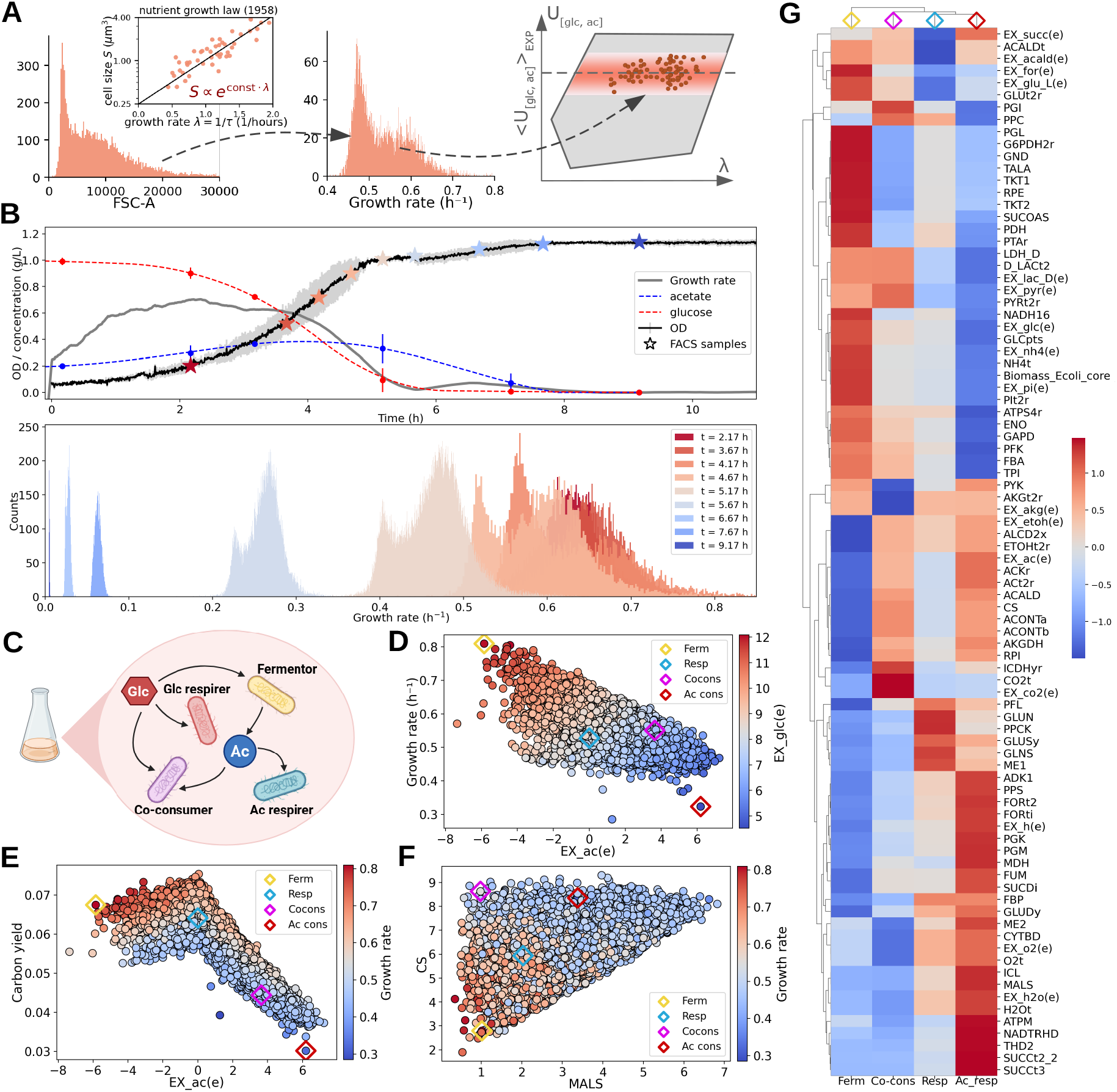
Single-cell metabolic flux inference of the diauxic shift in *E. coli*. **A**. Flow cytometry (FACS) data quantify single-cell sizes. Applying the exponential growth law yields single-cell growth rate distributions. Growth rates are fixed for each cell in the metabolic network model, and a maximum entropy approach fits the average glucose and acetate uptake rates using two independent Lagrange multipliers. **B**. Top: Experimental time courses from Figures 1 and 2, highlighting the FACS measurement time points. Bottom: Single-cell growth rate distributions inferred from FACS data at the corresponding time points. **C**. Schematic representation of distinct metabolic phenotypes coexisting within the population. **D, E, and F**. Scatter plots at *t* = 4.17 h (immediately preceding the diauxic shift): **D**. growth rate versus glucose and acetate fluxes; **E**. carbon yield versus acetate flux and growth rate; **F**. citrate synthase versus malate synthase flux and growth rate. Representative examples of the four metabolic phenotypes shown in **C** are highlighted in different colors. **G**. Clustermap of fluxes for all metabolic reactions in the core model across the four representative cells. Fluxes are Z-score normalized across cells; blue and red indicate low and high values, respectively.

The timing of the measurements relative to bulk culture dynamics, together with the single-cell growth-rate distributions, is shown in Fig. 3B. These distributions reveal substantial heterogeneity, particularly around the diauxic transition, consistent with the coexistence of multiple metabolic strategies schematically illustrated in Fig. 3C. Scatter plots at *t* = 4.17 h, immediately preceding the switch, show correlations between growth rate and glucose/acetate fluxes (Fig. 3D), carbon yield and acetate exchange (Fig. 3E), and TCA cycle fluxes (Fig. 3F). Four representative cells corresponding to the phenotypes in Fig. 3C are highlighted in distinct colors. Fig. 3G summarizes the metabolic variability across these cells as a clustermap of *Z*-score–normalized fluxes for all reactions in the core network.

From a computational standpoint, the inference problem might appear prohibitively high-dimensional (*N* × *N*_*c*_ ∼ 10^7^ effective degrees of freedom). However, all cells share the same flux polytope *P* and differ only through one individual inequality constraint on the growth rate, enabling efficient parallelization (Appendix D1), with Lagrange multipliers inferred via Boltzmann learning. Single-cell phenotypes span a continuous range from pure glucose consumption with acetate overflow to co-consumption, with almost no cells relying exclusively on acetate. These distributions are narrower than those from the population-level maximum entropy analysis, reflecting individual rather than population-average constraints.

Taken together, these results confirm that unbiased inference naturally points to a continuous, heterogeneous landscape of single-cell metabolic states characterized by mixed substrate utilization. Yet the Lagrange multipliers remain purely statistical constructs with no direct mechanistic interpretation. This motivates the complementary dynamical framework of the following section.

### Continuous consumer–resource model

The previous sections showed that unbiased inference points to a continuous, heterogeneous picture of single-cell metabolic states with mixed co-consumption rather than sharp transitions. Yet the Lagrange multipliers have no explanatory power—they are statistical devices, not mechanistic parameters. The consumer–resource picture of the first section provides a mechanistic explanation but within a framework too rigid to capture co-consumption quantitatively. Here we reconcile the two approaches by generalizing constraint-based consumer–resource models to continuous metabolic states [13].

To simplify the discussion we identify the metabolic state of a bacterium with the vector of its metabolic fluxes, *x* ≡ **v** = (*v*_1_, …, *v*_*n*_), and denote by *N*_*x*_ the density of bacteria in state *x*. The growth rate is determined by both the external environment *E* and the internal metabolic state *x*, λ = λ(*x, E*). We write a chemostat model allowing for random transitions between metabolic states. Denoting by *D* the dilution rate, by *C*_*i*_ (*i* = 1, …, *m*) the concentration of chemical *i* in the reactor, by *C*_*0,i*_ its concentration in the feed, and by *u*_*x,i*_ the uptake of chemical *i* at state *x* (negative values indicate secretion), the model reads

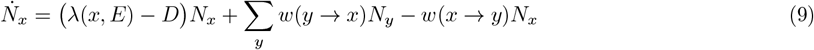

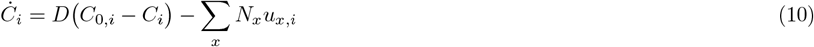

with *w*(*x* ⟶ *y*) the transition rate from state *x* to state *y*. Having identified *x* with the flux vector, Ω(*t*) is the polytope of allowed fluxes at time *t*, determined by stoichiometric and thermodynamic constraints that depend on the nutrient concentrations *C*_*i*_(*t*). The above system represents a large family of models, specified once w and the dependence of Ω(*t*) on the chemical densities are given. In the continuous limit, when *x* = (*v*_1_, …, *v*_*n*_) takes continuous values, many microscopic details of *w* (*x* ⟶ *y*) are integrated out. Defining the probability *p*_*x*_ = *N*_*x*_/*ρ* with *ρ* = Σ_*x*_ *N*_*x*_, under quite general assumptions [56, 57] the system (9,10) can be expanded to obtain a diffusion term [13] with effective diffusion constant 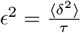, where *δ* is the typical magnitude of a single phenotypic jump (i.e. the change in flux vector upon a transition event) and τ is the characteristic time between successive jumps. Intracellular regulation further biases transitions towards higher-growth-rate configurations; this is encoded as a drift term whose potential coincides with the growth-rate bias of the MaxEnt ansatz. This regulatory advection gives rise to a drift towards higher λ [58], with strength controlled by a parameter *β* analogous to the MaxEnt Lagrange multiplier. Crucially, this bias does *not* act through replication (already captured by the replicator term) but mimics intracellular regulatory mechanisms pushing cells towards more efficient metabolic states. Denoting ⟨*u*⟩ = Σ_*x*_ *p*_*x*_*u*_*x*_, the continuous limit reads

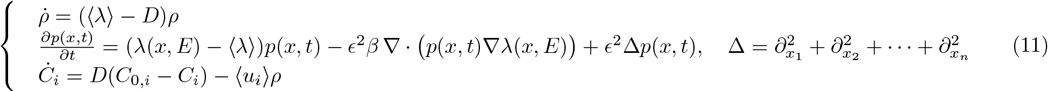

Where the second term, a Fokker–Planck-like equation, reduces to pure diffusion when *β* = 0, and the product *ϵ*^2^*β* realizes a fluctuation–dissipation-like relation between phenotypic noise and regulatory drift [58]. The polytope Ω(*t*) ℝ^*n*^ depends on the concentrations *C*_*i*_(*t*) and thus on time. To conserve total probability, the system must satisfy insulating boundary conditions,

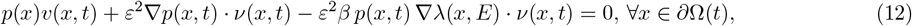

where 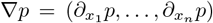 is the boundary of Ω(*t*),v is the outward unit normal, and v is the normal boundary velocity. This defines a moving-boundary problem for the trajectory and a free-boundary problem for the steady state [59], both to be solved self-consistently (see Appendix E1 for an explicitly solvable 1D toy model and Appendix E for the full multidimensional formulation). The resolution of such problems is computationally demanding; we therefore work with a simplified *E. coli* metabolic network.

Figure 4 presents the results of the continuous consumer–resource model applied to a simplified three-dimensional metabolic network of *E. coli* (Panel A), in which glucose uptake, acetate exchange, biomass production and respiratory flux capture the essential carbon-utilization trade-offs of the diauxic transition. Parameters are calibrated from the full constraint-based model (Appendix E 2), with best-fit dynamical values *β* ∼ 18 h and ϵ ∼ 0.6 mmol gDW^−1^ h^−1/2^.

**FIG. 4.**
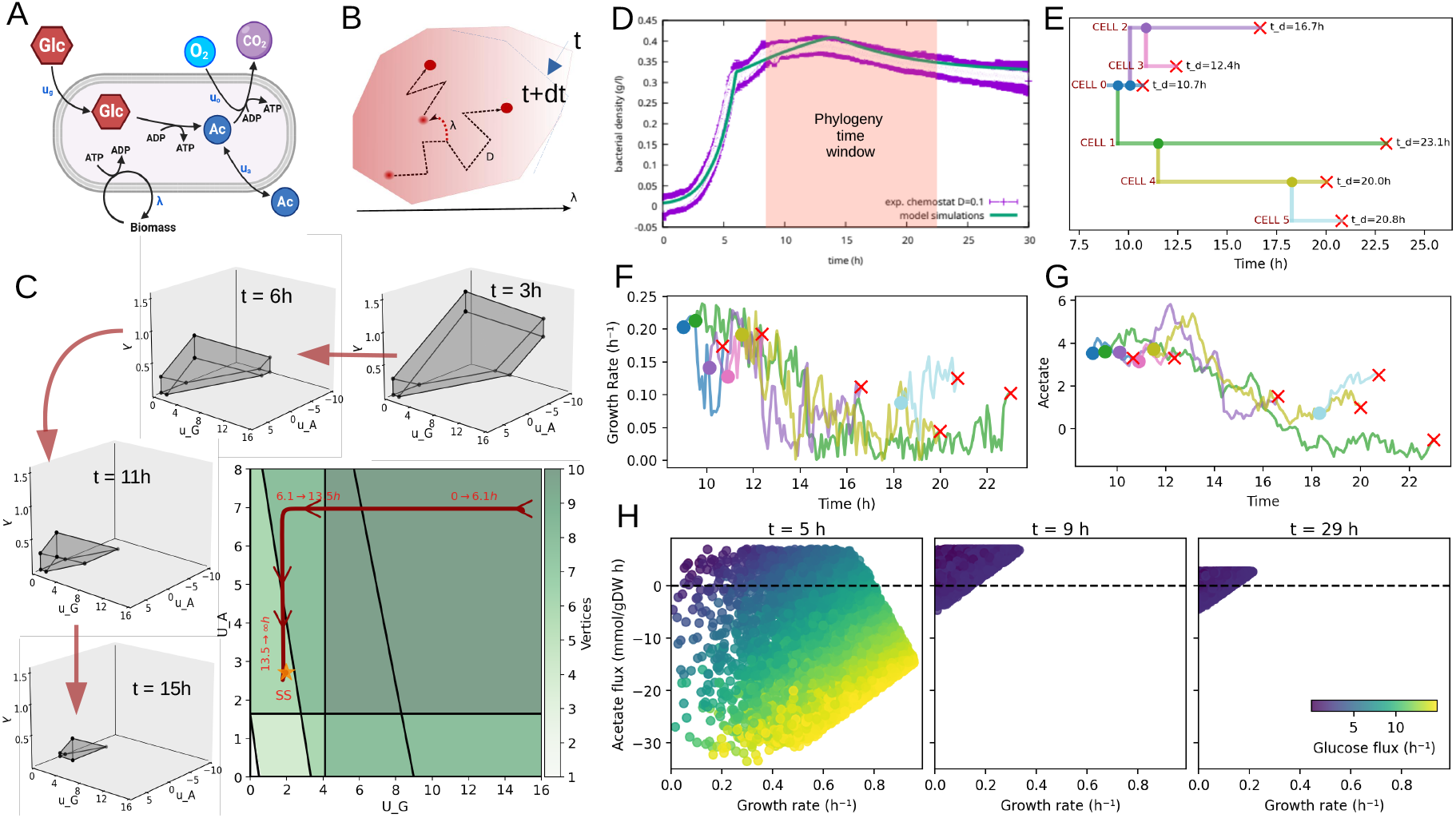
Network-based continuous consumer–resource model with moving metabolic boundary conditions. **A**. Simplified network model of E. coli metabolism. **B**. Sketch of diffusion–replication dynamics in the flux polytope with moving boundary conditions. **C**. Topological transformation of the flux polytope as a function of the upper bounds *U*_*G*_ and *U*_*A*_ on glucose and acetate uptake (which depend on the extracellular concentrations via Michaelis–Menten kinetics); the highlighted trajectory reconstructs the experimental trend. **D**. Comparison between model simulations and experimental data (bacterial density vs. time). Unlike the two-state discrete model, the continuous model reproduces complex dynamical trends, e.g. relaxation after a “second switch”. **E**. A randomly selected lineage (phylogenetic tree) showing birth and death times over 13 h. **F**,**G**. Single-cell acetate flux and growth rate for the lineage in **E. H**. Single-cell metabolic flux distributions (acetate flux vs. growth rate; glucose uptake: color code) at three time points (5, 9, and 29 h) spanning the switch.

Within this flux space, phenotypic diffusion, growth-rate-dependent replication, and the time-dependent deformation of the feasible polytope jointly shape the population’s metabolic landscape (Panel B). As nutrients are consumed, the polytope undergoes a sequence of topological transformations (Panel C; full classification in Appendix E 3): most strikingly, the acetate switch corresponds to a qualitative change in the geometry of the accessible metabolic space itself (the diffusionless limit is analyzed in Appendix E4). Fitting the two free parameters ϵ and β against experimental OD curves yields quantitative agreement with the growth dynamics across the entire diauxic transition—including the non-monotonic relaxation that eluded the two-state model (Panel D).

The model is simulated as a Moran process at fixed population size (Appendix E5; a 1D Crank–Nicolson benchmark is presented in Appendix E 1), producing lineage-resolved histories of individual cells. Panel E shows one such phylogenetic tree spanning thirteen hours, and Panels F and G trace the growth rate and acetate flux along its branches. What emerges is a picture of substantial intracellular dynamism: growth rates wander around the population mean and occasionally plunge well below it near the switch, while acetate fluxes reverse sign as cells drift between overflow and uptake. Daughters inherit their parent’s metabolic state at division, so that correlated runs of co-consumption or fermentation persist along lineage branches for a few generations before phenotypic diffusion erases the memory. The resulting single-cell flux distributions at representative time points (Panel H) recover the same continuous, co-consuming phenotypes identified by the maximum entropy inference—closing the loop between the statistical and dynamical descriptions and providing a dynamical account of the heterogeneity that both frameworks detect.

## DISCUSSION

Growth metabolism is a defining trait of microbial life, and understanding how metabolic strategies are distributed across individual cells is essential for interpreting population-level behavior, ecological interactions, and evolutionary adaptation. Here we presented a quantitative, multi-scale analysis of metabolic heterogeneity during one of the most classical yet still conceptually rich systems in microbiology: the glucose–acetate diauxic shift in *Escherichia coli*.

Recent experiments have challenged the textbook picture of diauxie as an abrupt, population-wide metabolic switch, revealing instead a robust regime of glucose and acetate co-consumption at the population level [5, 9, 10, 17]. The central question motivating this study was whether this coexistence reflects a mixture of specialized subpopulations or, rather, the emergence of intermediate metabolic states within individual cells. By combining consumer–resource modeling, constraint-based metabolic analysis, and maximum entropy inference, we provide converging evidence in favor of the latter scenario.

It is worth stressing that the glucose–acetate diauxie is mechanistically rather different from the paradigmatic glucose– lactose case underlying the textbook picture, and that this difference makes single-cell co-consumption *a priori* more plausible in our setting. Lactose utilization in *E. coli* relies on inducible enzymes encoded by the *lac* operon, whose expression is strongly repressed by glucose via cAMP–CRP-mediated catabolite repression and lac inducer exclusion [2, 3]. On a glucose+lactose mixture cells therefore operate in a near-binary regulatory regime that strongly biases towards an abrupt sequential switch. In contrast, acetate metabolism is supported by two routes with very different regulatory logic: the low-affinity, reversible Pta–AckA pathway, whose enzymes are essentially *constitutive* and active under most growth conditions, and the high-affinity, irreversible acetyl-CoA synthetase Acs, which is *inducible* and tightly controlled by cAMP–CRP [6, 9, 10]. The constitutive Pta–AckA arm operates in both directions depending on its thermodynamic driving force [9, 10, 17], so that the system already at the regulatory level admits a continuous range of mixed phenotypes—providing a natural mechanistic substrate for the heterogeneous co-consuming states recovered by our inference framework.

We first showed that a two-state consumer–resource model, in which cells optimally grow on either glucose or acetate, can reproduce many global features of batch and chemostat experiments. However, a constraint-based decomposition analysis demonstrated that observed co-consumption trajectories cannot in general be written as convex combinations of two optimally growing subpopulations, establishing a fundamental limitation of discrete, growth-optimizing population models and pointing to genuine single-cell co-consumption states enforced by the physicochemical constraints of the metabolic network.

Relaxing the assumption of strict optimality, we introduced a dynamical maximum entropy framework that infers time-dependent flux distributions directly from experimental growth data. This unbiased approach naturally predicts a continuum of metabolic phenotypes during the diauxic transition, spanning from glucose overflow to mixed glucose– acetate utilization. The co-consumption regime is not an epiphenomenon of population averaging, but a dominant and persistent feature of single-cell metabolism around the switch. The framework also predicts a rich exometabolome and broad flux variability, consistent with recent metabolomics and single-cell studies [15, 16, 43, 60].

By integrating flow cytometry measurements, we extended this approach to perform single-cell metabolic flux inference, reconstructing intracellular flux distributions conditioned on individually inferred growth rates and revealing correlated variations in uptake strategies, energy metabolism, and biomass production. We stress, however, that single-cell growth rates were not measured directly but inferred from the time evolution of forward-scatter (FSC) signals, used as a proxy for cell size under the assumption of exponential single-cell growth. This introduces caveats worth keeping in mind: FSC depends on optical properties beyond cell volume [61] and may be affected by cell shape, refractive index, and intracellular composition; the exponential-growth assumption may not hold around the diauxic switch, where cells may transiently slow down or arrest [14, 60]; and snapshot cytometry mixes cells at different cell-cycle positions. With these caveats in mind, these results reinforce the view that phenotypic heterogeneity during diauxic growth is best described as a continuous distribution over metabolic states.

Finally, we reconciled the statistical and dynamical perspectives by formulating a continuous consumer–resource model over metabolic state space. The population is described by a probability density over the flux polytope Ω(t) obeying a Fokker–Planck-like equation with three explicit ingredients: a replicator term encoding selection at the phenotypic level; a regulatory drift towards higher growth rates [58], mimicking intracellular regulation; and an isotropic diffusion term capturing unbiased phenotypic change. The polytope deforms in time through Michaelis–Menten couplings to the chemostat ODEs, turning the steady-state problem into a free-boundary problem and the trajectories into moving-boundary problems [59]. The two fitted parameter ϵ and β simultaneously determines the width of predicted single-cell flux distributions, the timescale of polytope adaptation, and the lineage-resolved fluctuations in Fig. 4, connecting classical chemostat theory [18, 19], replicator dynamics [13, 62], and constraint-based modeling [21, 22] into a unified, quantitatively predictive framework.

Taken together, this study suggests that even paradigmatic systems such as the *E. coli* diauxic shift conceal a rich landscape of single-cell metabolic states. Beyond their fundamental interest, results of this kind have direct practical implications: most industrial *E. coli* bioprocesses are run on glucose with significant acetate accumulation, and the population-level heterogeneity we recover is precisely the variability shown to control the robustness, productivity, and reproducibility of large-scale fermentations [12, 63–65]. From a biological standpoint, the existence of long-lived co-consuming and slow-growing single-cell states around the switch fits naturally into the bet-hedging interpretation of phenotypic heterogeneity [11, 65, 66]: when nutrient depletion is unpredictable, populations maintaining a continuous spread of metabolic strategies may outperform those locked into a single growth-optimal phenotype. Our framework does not by itself test the evolutionary hypothesis of bet-hedging, but it provides the natural quantitative language in which it can be formulated. Capturing this landscape requires moving beyond discrete switches and strict optimality, toward probabilistic and dynamical descriptions that embrace heterogeneity as a fundamental feature of living systems. These results resonate with a growing body of literature emphasizing suboptimality, heterogeneity, and metabolic flexibility as central organizing principles of microbial physiology [11, 12, 26, 28].

Several concrete directions emerge from this work. On the experimental side, time-resolved single-cell measurements simultaneously reporting substrate uptake, reporter gene expression, and intracellular metabolite levels—through, for example, microfluidic lineage tracking combined with fluorescent biosensors—would allow direct validation of the inferred flux distributions and co-consumption phenotypes [14, 16, 60]. Isotope-resolved metabolomics at the single-cell level would provide an independent handle on carbon routing through the TCA cycle and the Pta–AckA pathway [9, 15]. We note that the single-cell flux inference presented here is indirect, in that it projects cytometry-derived growth rates onto the metabolic polytope via maximum entropy; it does not measure intracellular fluxes per se. A natural next step would be to combine our framework with ^13^C isotope tracing and high-resolution mass spectrometry—the established tools of metabolic flux analysis at the population level [67–69]—to move towards a true single-cell flux analysis in which isotopomer data, rather than growth-rate proxies alone, constrain the intracellular flux map of each individual cell. A complementary test would be to apply our framework to *E. coli* mutants with perturbed Pta– AckA/Acs balance (e.g. ∆*acs*, ∆*ackA*, ∆*pta*), which display markedly altered co-consumption phenotypes [9, 10, 17]: our continuous formulation predicts how the entire single-cell flux distribution should shift in each genetic background, providing many simultaneous constraints. More broadly, the same framework applies to other diauxic shifts in *E. coli* and to growth on multiple carbon sources in non-model organisms, and is naturally suited to ecologically realistic settings in which bacteria face complex carbon mixtures [5], including spatially structured communities [70, 71].

On the theoretical side, incorporating enzyme allocation constraints and proteome-aware bounds into the maximum entropy framework would link metabolic heterogeneity explicitly to translational noise and ribosome partitioning, connecting our approach to coarse-grained growth laws [4, 8, 30]. Extending the continuous consumer–resource model to multi-species communities—where cross-feeding metabolites couple multiple strains—would address questions about metabolic niche partitioning and coexistence [45, 70, 72, 73]. More broadly, the moving-boundary formulation applies naturally to any system in which the feasible phenotypic space changes with environmental feedback, including catabolite repression in complex carbon mixtures [3, 5], osmotic or oxidative stress responses, and evolutionary adaptation in fluctuating environments [12, 62]. We anticipate that this combination of maximum entropy inference, constraint-based metabolism, and continuous consumer–resource theory will serve as a broadly applicable toolkit for quantifying and interpreting phenotypic heterogeneity in microbial physiology.

## METHODS

### Experimental methods

#### Strain handling

*E. coli* MG1655 experiments were initiated by inoculating a single colony into an LB pre-culture. After 18 hours, cells were transferred to M9 pre-cultures supplemented with 0.4% glucose. Batch and continuous growth experiments were carried out in M9 medium with a total volume of 25 mL using a 16-vessel bioreactor system (eVOLVER, FinchBio, USA), equipped with aeration to prevent oxygen limitation (Dan, manuscript in preparation). All experiments were carried out at 37 °C and medium stirring.

For batch experiments, M9 medium was supplemented with 1 g L^−1^ glucose and 0.2 g L^−1^ acetate; this is the reference condition used to fit the two-state consumer-resource and dynamical maximum entropy models in the main text. Additional batch experiments at different glucose and acetate concentrations were also performed and used as independent test conditions for the fitted models. Chemostat experiments included an initial batch phase (1 g L^−1^ glucose and equimolar 0.4 g L^−1^ acetate), followed by a continuous phase at dilution rates of 0.1 or 0.3 h^−1^. Pulse experiments were initiated from a batch phase supplemented only with glucose (0.3 g/L). After 23 hours, the feed was switched to glucose (1.3 g L^−1^). In a separate condition, cultures were supplemented with 0.5 g L^−1^ glucose, and an acetate pulse (1 g L^−1^) was added to the reactors after 11 hours.

### Analytics

Aliquots of 200 µL culture were taken at regular intervals. Half of each sample was filtered (0.2 µm) for metabolite analysis by chromatography (Column: BioRad Aminex HPX-87C; isocratic elution with 0.5 mM H_2_SO_4_ at 0.6 mL min^−1^, Agilent 1100 series, RI detector). The remaining half was immediately mixed with the styryl pyridinium dye RH414 (3 µM, MCE) to distinguish small debris from cells and analyzed by flow cytometry (Sony SH800S Cell Sorter), where forward scatter (FSC) was recorded as a proxy for cell size.

### Computational methods

#### Instantaneous growth rate estimation

Optical density (OD) measurements from replicate experiments were averaged to obtain mean growth curves and associated standard deviations. OD values were scaled (factor 0.35) and log-transformed to approximate cell density. The resulting signal was smoothed using a moving average filter (window size a = 100 time points) to reduce noise. Instantaneous growth rates were then computed as the numerical derivative of the smoothed log-transformed signal using finite differences, followed by an additional smoothing step.

#### Two-state consumer-resource model

To interpret the observed growth dynamics, we employed a two-state consumer–resource model describing *E. coli* populations growing on glucose and acetate (see Appendix A for full model formulation and Appendix A for parameter estimation details). The model considers two subpopulations consuming either glucose or acetate, with growth rates governed by Michaelis–Menten kinetics and yield coefficients. Cells can switch between metabolic states depending on nutrient availability, with transition rates modulated by glucose and acetate concentrations. The model additionally accounts for overflow metabolism, whereby excess glucose uptake leads to acetate secretion above a threshold uptake rate. Nutrient concentrations are dynamically coupled to population growth through uptake and dilution terms (in chemostat conditions). Model parameters were estimated by fitting to batch culture data using a simulated annealing optimization procedure (see Appendix A), and the resulting parameter set was subsequently used to predict dynamics in chemostat experiments. System behavior and stability were analyzed by computing steady states and performing a linear stability analysis of the resulting dynamical system (Appendix A).

#### Metabolic model and Flux Balance Analysis

All simulations were performed using the COBRApy (Constraint-Based Reconstruction and Analysis) toolbox with a modified version of the *Escherichia coli* core metabolic model [74], including the ACS reaction. Flux Balance Analysis (FBA) was used to compute steady-state flux distributions by maximizing the biomass objective function subject to stoichiometric and flux constraints (S*v* = 0, *v*_*min*_ ≤ v ≤ *v*_*max*_), using the GLPK linear programming solver. To characterize the feasible metabolic space, biomass production was systematically constrained across a range of values from zero to its maximum, and exchange fluxes were computed at each point. This procedure enabled reconstruction of the feasible flux envelope under growth constraints, which was used to generate the flux space representation shown in Figure 1D.

#### Dynamic maximum entropy

We developed a dynMaxEnt framework to infer time-resolved *E. coli* metabolic flux distributions under changing extracellular conditions by combining constraint-based modeling with entropy-weighted stochastic sampling (Appendix C). At each time point, fluxes v constrained by S*v* = 0 and bounds l ≤ *v* ≤ *u* were sampled with ACHRLto approximate the uniform distribution P_0_(*v*). This was reweighted as 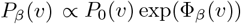, where 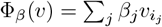 using a Metropolis–Hastings acceptance step min (1, *e*^∆Φ^). Bias coefficients were standardized by pilot-run flux variances and optionally annealed to improve mixing. Extracellular concentrations were updated via Michaelis–Menten kinetics, and biomass evolved according to sampled growth rates. Iterating across time points produced coupled trajectories of environmental conditions and flux distributions consistent with stoichiometric constraints and experimental growth dynamics.

## ACKNOWLEDGMENTS

D.D.M. acknowledges financial support from the grants PIBA 2024 1 0016 (Basque Government) and Project PID2023-146408NB-I00 funded by MICIU/AEI/10.13039/501100011033 and by FEDER, UE. A.F.F. acknowledges support from the Predoctoral Training Program for Non-Doctoral Research Personnel of the Basque Government’s Department of Education. D.M. acknowledges financial support from the grant UID/PRR/00208/2025 (https://doi.org/10.54499/UID/PRR/00208/2025) funded by Funda ç ão para a Ci ê ncia e a Tecnologia (FCT) and União Europeia - Estrutura de Missão Recuperar Portugal (UE - EMRP), and from the grant UID/00208/2025 funded by Fundação para a Ciência e a Tecnologia (FCT). D.M. expresses his gratitude to the Theoretical/Protocell Biophysics Laboratory of Biofisika Institute (CSIC, UPV/EHU) for the warm hospitality. K.S.K. was supported by NIH/NIGMS Grant No. 1R01GM138530-01. D.S. acknowledges support by the U.S. Department of Energy, Office of Science, Office of Biological & Environmental Research through the Microbial Community Analysis and Functional Evaluation in Soils SFA Program (m-CAFEs) under contract number DE-AC02-05CH11231 to Lawrence Berkeley National Laboratory and the National Science Foundation (grants NSFOCE-BSF 1635070 and the NSF Center for Chemical Currencies of a Microbial Planet-C-CoMP article #089). D.S., L.K., I.D. were partially supported by grant 1R21CA279630-01 from the National Cancer Institute at the National Institutes of Health.

## Appendix A: Two-state discrete consumer–resource model

Consider a population of *E. coli* cells growing in a chemostat with dilution rate *D* (the case of batch can be recovered setting *D = 0*). Cells can be either consuming glucose (*N*_*G*_, *G*) or acetate (*N*_*A*_, *A*). Cells can shift from one to the other population at a certain rate (*W* ). The variation in the size of each population and the nutrient concentration is then given by:

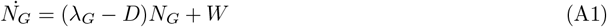

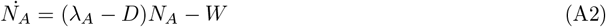

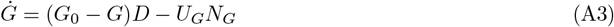

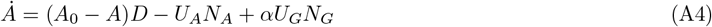

where *A*_0_, *G*_0_ stand for the concentration level of acetate and glucose in the feed. Cells within each population grow at a rate λ that is a function of Michaelis Menten kinetics (where *V*_*max*_ and *K* are the maximum velocity and the Michelis Menten constant) and the yield (γ) for each compound:

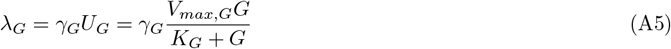

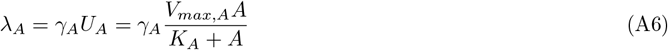

Additionally, the glucose yield depends on the kind of metabolism that glucose consumers perform. Above a certain threshold (*U*_*G,t*_) that we refer to as the overflow constraint following [8], cells start fermenting part of the glucose into acetate so that equation A15 can take two different forms:

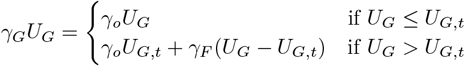

Following this same reasoning, the parameter α in equation A4 reflects the amount of acetate produced by the fermenting cells as follows:

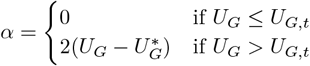

where the factor 2 corresponds to carbon stoichiometry from glucose to acetate in the fermentation pathway. Finally, we introduced the term W to refer to the transition between populations, that is further extended as:

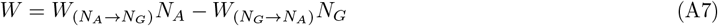

where 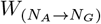 and 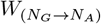 correspond to the transition rate from acetate to glucose consumers and the other way around respectively and have the following form:

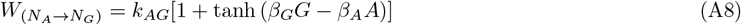

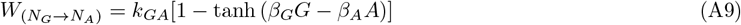

This leaves us with a model of 4 species (*N*_*G*_, *N*_*A*_, *G* and *A*), 3 control parameters (*D, G*_*0*_ and *A*_*0*_) and 12 constant parameters.

### Parameter Estimation and Model Fitting

Model parameters were estimated by fitting a consumer–resource model to experimental measurements of biomass (optical density, OD) and extracellular metabolite concentrations (glucose and acetate). Parameter inference was performed using a stochastic global optimization approach based on simulated annealing. Two distinct datasets were used:

1. **Fitting dataset (batch experiments):** Time-resolved measurements of optical density and extracellular metabolite concentrations obtained from batch culture experiments were used to infer model parameters.
2. **Test dataset (chemostat experiments):** Independently collected chemostat experiments were used to evaluate the predictive capability of the model using the parameter set obtained from batch data fitting. These data were not used during parameter estimation.

Model outputs used for the fitting included total biomass, acetate concentration, and glucose concentration. Parameter estimation was performed by minimizing a weighted sum of squared residuals between model predictions and experimental data. The objective function was defined as:

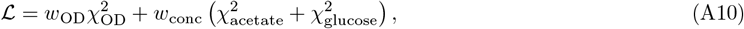

where each term corresponds to a weighted least-squares error:

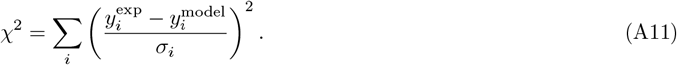

Here, 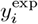 and 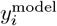 denote experimental and simulated values, respectively, and *σ*_*i*_ is the experimental standard deviation. Optical density was assigned a higher weight (*w*_OD_ = 10) relative to metabolite concentrations (*w*_conc_ = 1) to prioritize accurate fitting of biomass dynamics due to the nature of experimental measurements (continuous OD and discrete concentrations). Model parameters were constrained within biologically plausible bounds during optimization. Additionally, weak quadratic priors were imposed on a subset of parameters to penalize deviations from their initial estimates. This regularization term was added to the objective function during optimization. Global optimization was performed using a simulated annealing algorithm. Starting from an initial parameter set, new candidate parameter vectors were generated through stochastic perturbations. For most parameters, multiplicative (log-scale) perturbations were applied:

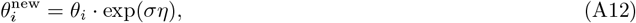

where *η∼ 𝒩* (0, 1) and *σ* is a step-size parameter. This approach ensures scale-invariant exploration across parameters spanning multiple orders of magnitude. Two parameters were held fixed during optimization. Candidate solutions were accepted according to the Metropolis criterion:

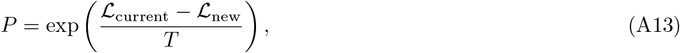

where *T* is a temperature parameter controlling acceptance of uphill moves. The temperature was gradually decreased according to an exponential cooling schedule:

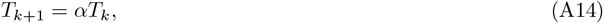

The perturbation scale was also gradually reduced during optimization to allow coarse exploration at early stages and fine-tuning at later stages. The best parameter set encountered during the annealing trajectory was retained as the final estimate, which is shown in Table I.

**TABLE I.** Parameter values.

| Parameter | Value | Parameter | Value |
| --- | --- | --- | --- |
| $K_{m,G}$ | $0.0043 \text{ h}^{-1}$ | $Y_{G,ferm}$ | 0.024 |
| $V_{max,G}$ | $12.014 \text{ mmol}/(\text{gDWh})$ | $U_{overflow}$ | $6.98 \text{ mmol}/(\text{gDWh})$ |
| $K_{m,A}$ | $0.0696 \text{ h}^{-1}$ | $\beta_G$ | 5.59 |
| $V_{max,A}$ | $7.98 \text{ mmol}/(\text{gDWh})$ | $\beta_A$ | 0.010 |
| $Y_A$ | 0.008 | $k_{GA}$ | 8.08 |
| $Y_{G,resp}$ | 0.08 | $k_{AG}$ | 0.02 |

The optimized parameter set obtained from batch experiments was subsequently used without further adjustment to simulate chemostat experiments. Model predictions were compared against the independent test dataset to assess predictive performance and generalizability.

### Validation and limitations of the two-state consumer–resource model across dynamical regimes

To further assess the validity of the two-state discrete consumer–resource model introduced in the main text, we compared its predictions against experimental data across a range of dynamical conditions, including steady-state chemostat growth, nutrient shifts, and pulsed perturbations. The results are summarized in Fig. 5.

**FIG. 5.**
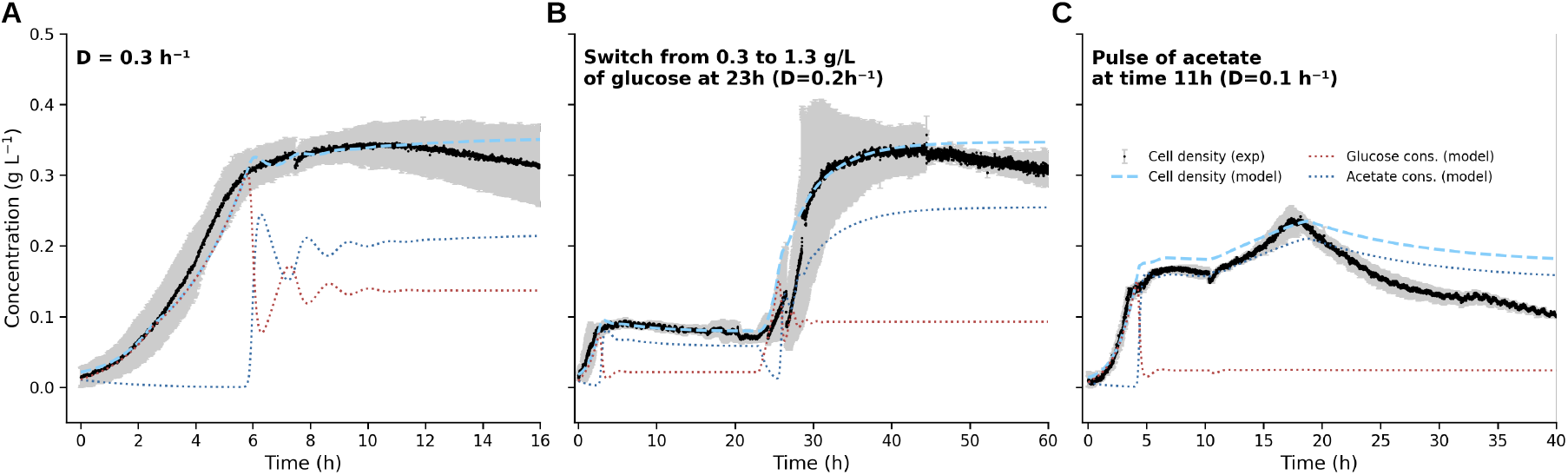
Consumer–resource model predictions for chemostat experiments. **A**. Continuous culture at a dilution rate of 0.3, h^−1^ with feed concentrations of 1 g/L glucose and 0.4 g/L acetate. **B**. Continuous culture at a dilution rate of 0.2, h^−1^, where the glucose feed concentration was increased from 0.3 g/L to 1.3 g/L at 23 h. **C**. Pulse experiment at a dilution rate of 0.1, h^−1^ with a constant glucose feed of 0.5 g/L; acetate was added as a single pulse at 11 h with a concentration of 1 g/L.

At a dilution rate *D* = 0.3 h^−1^ (Fig. 5A), the model reproduces the overall dynamics of biomass accumulation with good quantitative agreement. However, small deviations emerge during the growth phase and at longer times, where the model is not able to capture the decay observed in the experiments, similarly to Fig. 1C.

We next consider a step increase in glucose concentration from 0.3 to 1.3 g L^−1^ at *t* = 23 h, under a chemostat regime with *D* = 0.2 h^−1^ (Fig. 5B). The model successfully predicts the qualitative response, including the rapid increase in biomass and the transient metabolic adjustment. The sharp transition is driven by the fast switching rate from acetate-to glucose-consuming subpopulations encoded in the model. Quantitatively, however, it appears to miss the lag in the transition observed experimentally, and again it does not capture the final decay.

A more stringent test is provided by an acetate pulse at *t* = 11 h (Fig. 5C) under low dilution (*D* = 0.1 h^−1^). In this case, the experimental trajectory displays a pronounced overshoot in biomass followed by a slow decay toward steady state. While the model captures the immediate response to the acetate addition, it fails to reproduce the non-monotonic relaxation dynamics, instead predicting a smoother and more gradual return to equilibrium.

These observations support the conclusion drawn in the main text: while the discrete two-subpopulation picture serves as a useful baseline, it is insufficient to fully capture the dynamics of co-consumption and metabolic adaptation. This motivates the development of a continuous framework in which cells can explore a spectrum of metabolic states, as introduced in the following sections.

### Linear stability analysis

We performed a local stability analysis of the steady state of the four-dimensional consumer–resource system defined in Eqs. (A4–A15) by evaluating the Jacobian matrix at steady state and computing its eigenvalues and eigenvectors as a function of the dilution rate *D*. This analysis provides insight into both the stability of the steady state and the dominant dynamical modes governing perturbations. To find equilibrium points, we set all derivatives to zero:

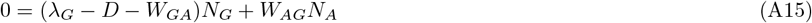

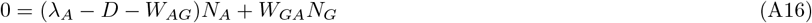

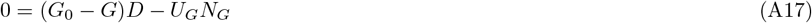

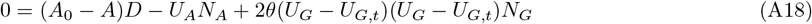

Where the *θ* function is either 1 or 0 depending on whether there is the overflow or not. Solving these equations provides the equilibrium point (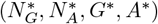, *G*^*^, *A*^*^). We perform a linear stability analysis by linearizing the system around the equilibrium point. Define small perturbations:

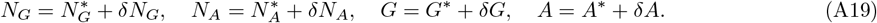

Expanding the system to first order in perturbations, we obtain the Jacobian matrix (J):

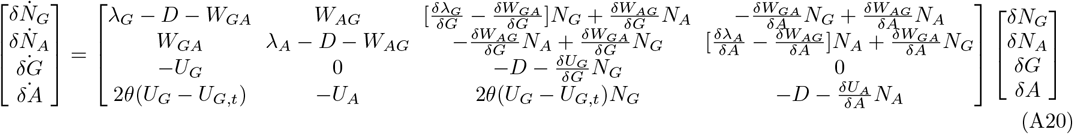

The stability of the system is then determined by the eigenvalues of the Jacobian matrix that are found by solving:

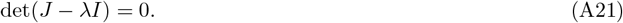

The real part of each eigenvalue, 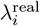, determines the stability of the corresponding mode. Negative real parts indicate that perturbations decay and the steady state is locally stable, whereas positive real parts indicate instability. The presence of a non-zero imaginary part, 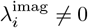, signals oscillatory dynamics and is indicative of complex conjugate eigenvalue pairs. The associated eigenvectors provide information about which state variables dominate each mode. In particular, the relative magnitudes of the components corresponding to *N*_*G*_, *N*_*A*_, *G*, and *A* indicate whether a given dynamical mode is primarily driven by population dynamics or resource concentrations.

### Structure of the dynamical modes

The spectrum of the Jacobian reveals four qualitatively distinct modes. The first eigenmode exhibits a real eigenvalue that increases with the dilution rate and eventually becomes positive. This mode is non-oscillatory and transitions from a mixed contribution of variables at low *D* to being dominated by the acetate concentration *A* at higher *D*. This behavior indicates the emergence of an instability associated with acetate-mediated feedback. Biologically, this reflects the increasing importance of overflow metabolism and acetate accumulation as the dilution rate increases.

The second eigenmode displays a transition from purely real to complex eigenvalues as *D* increases, with a non-zero imaginary component emerging beyond a critical dilution rate. This indicates the onset of oscillatory dynamics through a Hopf-like mechanism. The corresponding eigenvector shifts from being dominated by glucose-related variables to a stronger coupling with acetate dynamics, suggesting that the oscillations arise from delayed feedback between glucose consumption, acetate production, and population switching.

The third eigenmode remains unstable over a broad range of dilution rates, with a positive real part and a growing imaginary component. Notably, this mode exhibits a sharp transition in its eigenvector composition at an intermediate dilution rate. This transition coincides with a collapse of the acetate component and a redistribution of weight across the remaining variables. This behavior is consistent with a nonlinear regime shift driven by the switching term *W* and the activation of overflow metabolism. It suggests the presence of bistability or rapid transitions between glucose-dominated and acetate-dominated growth regimes.

The fourth eigenmode is characterized by a large negative real eigenvalue and no imaginary component, indicating a fast and strongly stable mode. Its eigenvector is almost entirely aligned with the acetate concentration, implying that this mode corresponds to rapid equilibration of acetate.

### Global dynamical behavior

Taken together, the eigenvalue spectrum shows that steady-state stability is strongly controlled by the dilution rate, with increasing *D* eventually driving at least one eigenvalue positive and thus destabilizing coexistence, while the appearance of complex eigenvalues indicates the possibility of oscillatory dynamics. A key transition occurs at intermediate dilution rates (*D* ≈ 0.2–0.3), where both eigenvalues and eigenvectors change abruptly, marking the onset of overflow metabolism in which glucose consumption induces acetate production and reorganizes system feedbacks. Overall, the system dynamics emerge from the coupled effects of metabolic feedback via acetate, nonlinear phenotype switching mediated by *W*, and resource-dependent growth kinetics, whose interaction introduces delayed cross-population coupling and drives instabilities, oscillations, and regime shifts.

## Appendix B: Experimental trajectory in the polytope

We extended the analysis of points 1–5 in Figure 1B and D to evaluate the continuous trajectories using a smoothed fit of the curves (Fig. 7A). Uptake fluxes and growth rates were then computed from concentration and OD curves and projected onto the feasible region of the model in Fig. 7B. The trajectory initially lies in the acetate production region associated with high glucose uptake, then moves through a transitional regime toward the co-consumption boundary, before approaching states consistent with acetate utilization.

**FIG. 6.**
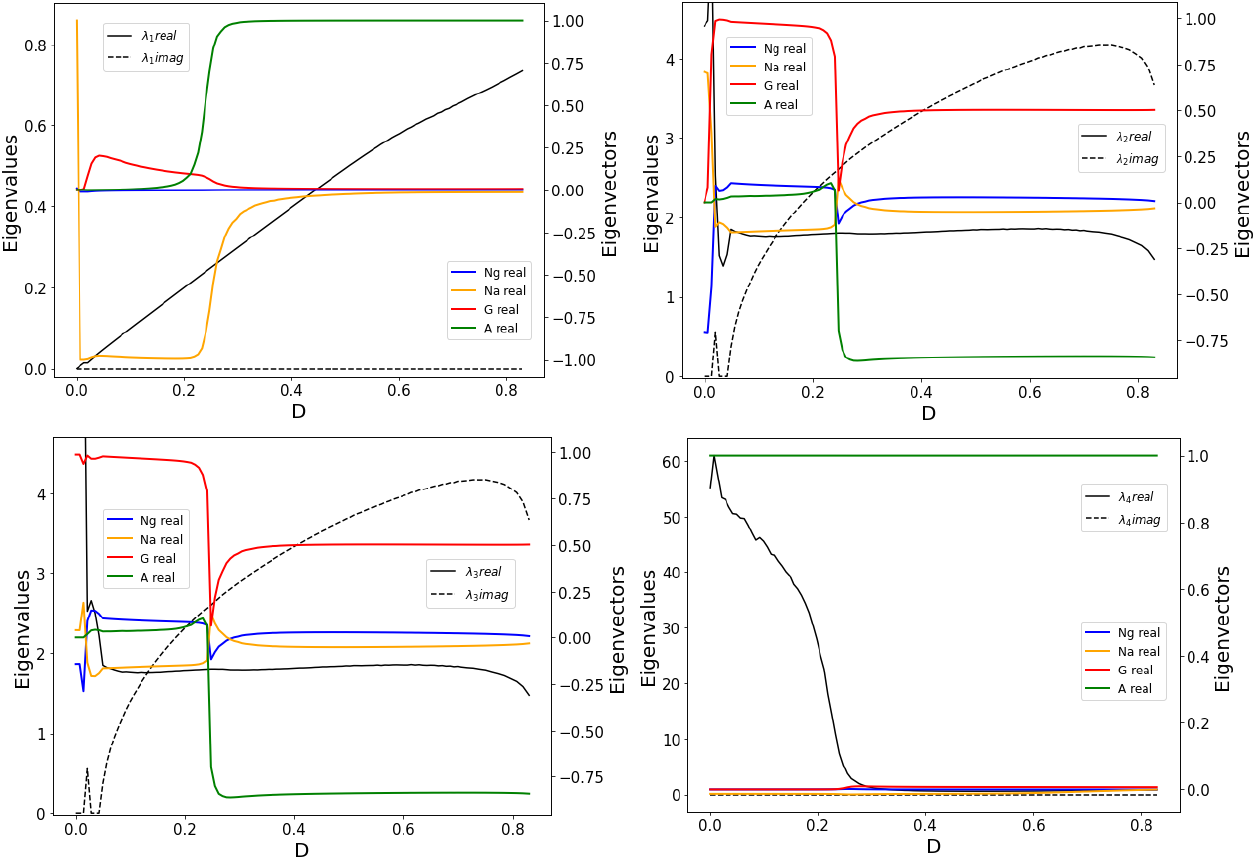
Eigenvalues and eigenvectors of the model as a function of the dilution rate for a medium composition of 1g/L glucose + 0.4g/L acetate

**FIG. 7.**
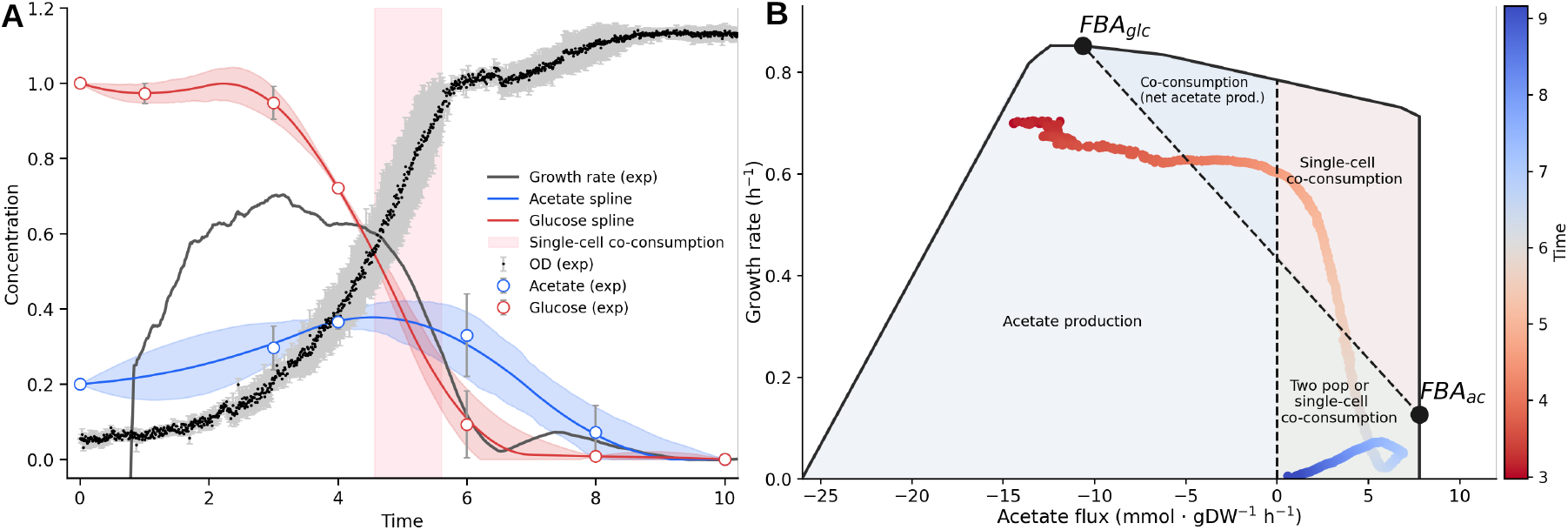
Experimental trajectory of growth rate and acetate consumption. **A**. Smoothed fits of experimental acetate and glucose concentrations together with measured cell density. The highlighted window corresponds to a phase of obligate single-cell co-consumption. **B**. Polytope of the core metabolic model with the experimental trajectory of growth rate versus acetate uptake projected from A. The color gradient indicates time progression.

The time-resolved coloring shows that single-cell co-consumption is the only regime that can describe the experimental data within the highlighted time window in panel A (4.57 to 5.6 hours). This interval indicates that co-consumption begins approximately one hour before the diauxic shift, supporting the idea that this phenomenon reflects a gradual change in metabolic diversity rather than an abrupt transition between two distinct metabolic states.

## Appendix C: Dynamic maximum entropy framework for flux distribution inference

We developed a dynamic maximum entropy (dynMaxEnt) framework to infer time-dependent distributions of metabolic fluxes in *Escherichia coli* under varying extracellular conditions. The approach integrates constraint-based modeling with stochastic sampling and an entropy-based reweighting scheme. Constraint-based metabolic models describe feasible steady-state flux vectors *v* ∈ ℝ^*n*^ that satisfy the stoichiometric constraints

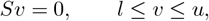

where *S* is the stoichiometric matrix and *l, u* are lower and upper flux bounds. The feasible region ℱ forms a convex polytope. Standard sampling methods such as ACHR or OptGP approximate the uniform distribution over ℱ, denoted *P*_0_(*v*). Sometimes, however, we are interested in *biased* sampling that favours flux distributions exhibiting large activity through selected reactions. For instance, one might want to draw samples with elevated ATP maintenance or glucose uptake while maintaining feasibility. To bias the distribution we introduce a scalar score (or negative energy) function Φ(*v*) and define the Boltzmann-weighted target distribution

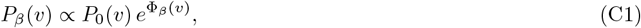

where Φ_*β*_(*v*) encodes preferences through a set of coefficients *β* = (*β*_1_, …, *β*_*k*_) corresponding to target reactions *T* = {*i*_1_, …, *i*_*k*_}. A simple and effective choice is a weighted linear combination of fluxes:

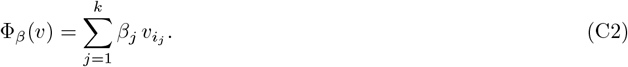

Each coefficient *β*_*j*_ controls the bias strength for reaction *i*_*j*_. Larger *β*_*j*_ values give stronger preference for higher flux through that reaction. Setting all *β*_*j*_ = 0 recovers the uniform sampler.

### 1. Sampling algorithm

The BiasedSampler modifies the standard Artificial Centering Hit–and–Run (ACHR) procedure used in COBRApy. The key change is to introduce a Metropolis–Hastings acceptance step that enforces the Boltzmann weight (C1).

#### Algorithm 1

Metropolis–Hastings biased ACHR step

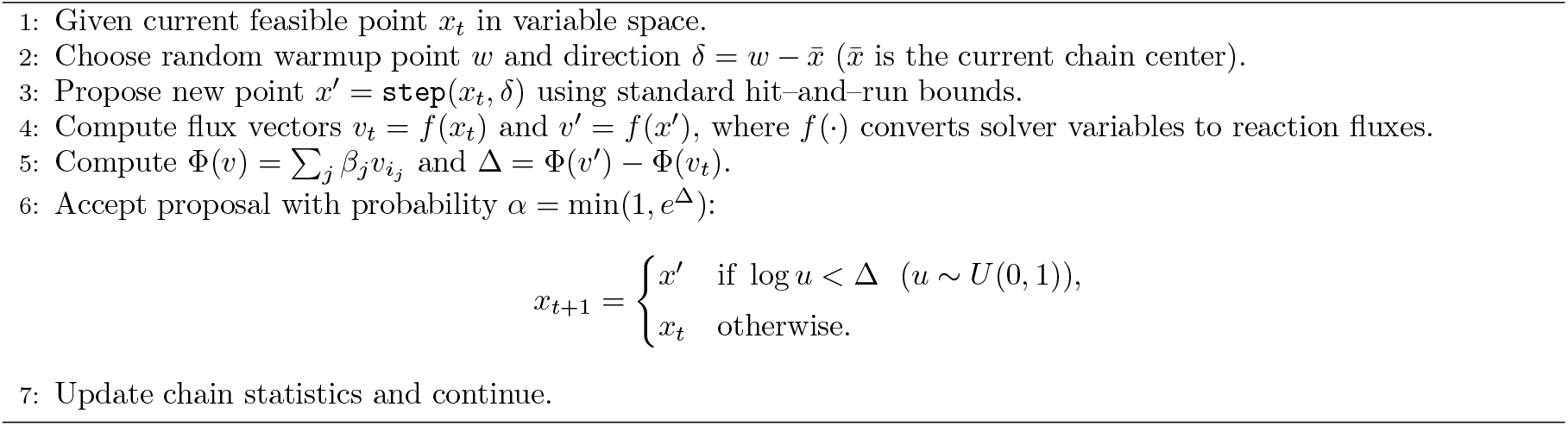

Using the log form of the acceptance test avoids numerical overflow for large Φ differences.

### Automatic scaling of *β* values

Fluxes can have different natural magnitudes. To prevent one reaction from dominating solely because its fluxes are large in absolute terms, the implementation offers an *automatic scaling* option:

1. Run a short unbiased pilot sampling of length *N*_pilot_.
2. Estimate the standard deviation *σ*_*j*_ of each target reaction.
3. Replace user-provided raw coefficients by

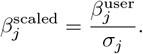

This standardization ensures that all targets contribute on a comparable scale.

### Annealing of *β*

To improve mixing, the sampler can *anneal* the bias strength over the first *N*_anneal_ iterations:

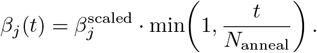

Early iterations are thus almost unbiased and gradually become fully biased.

### Calibration of *β*_*λ*_ to match growth rate

The biasing parameter *β*_*λ*_ associated with the biomass reaction was determined iteratively at each simulation time point in order to match the experimentally measured instantaneous growth rate. For a given target growth rate ⟨*g*⟩, an initial estimate of *β*_*λ*_ was obtained either from a precomputed lookup table of reference simulations (by minimizing the normalized deviation between predicted and target growth) or, for subsequent time points, by perturbing the previously accepted value. Using this candidate *β*_*λ*_, flux distributions were sampled with the Boltzmann-biased ACHR algorithm, and the resulting mean biomass flux ⟨*v*_biomass_⟩ was compared to ⟨*g*⟩. The relative error was then used to update *β*_*λ*_ in a stochastic, adaptive manner, with step sizes proportional to both the magnitude and sign of the error. Updates that reduced the error function (which also penalized deviations from previous *β*-estimates to ensure temporal smoothness) were accepted. This procedure was repeated until the sampled flux ensemble was consistent with the imposed instantaneous growth constraint.

### Dynamic Update of Extracellular Conditions

At each time step, extracellular glucose and acetate concentrations were updated based on the sampled exchange fluxes. Uptake rates were modeled using Michaelis–Menten kinetics, with parameters calibrated for glucose (*K*_*g*_, *V*_*g*_) and acetate (*K*_*a*_, *V*_*a*_). The biomass density *N* was propagated exponentially according to the sampled growth rate. These updates defined the dynamic boundary conditions of the metabolic model.

### Iterative Simulation

The above procedure was repeated across experimental time points. At each iteration:

1. The metabolic model was updated with current extracellular concentrations.
2. *β* was estimated to match the experimental average growth rate
3. The distribution of metabolic fluxes was sampled with the estimated *β*.
4. Extracellular metabolite concentrations and biomass density were updated.

This iterative scheme generates time-resolved maximum entropy distributions of metabolic fluxes that are consistent with both stoichiometric constraints and experimental growth measurements.

The code implementing both the CR model and the dynamic maximum entropy (dynMaxEnt) framework is available in the public GitHub repository:

https://github.com/arianferrero/dynMaxEnt.git.

The dynMaxEnt framework accurately captures the temporal evolution of growth dynamics across metabolic regimes (Fig. 8A). In contrast to standard FBA, which predicts near-constant maximal growth capacity, the inferred growth rate closely follows the experimental measurements, including the sharp decline associated with the metabolic transition. This transition is accompanied by a pronounced increase in the inferred bias parameter *β*_*λ*_, indicating a rapid reweighting of flux distributions to enforce the experimentally observed slowdown.

**FIG. 8.**
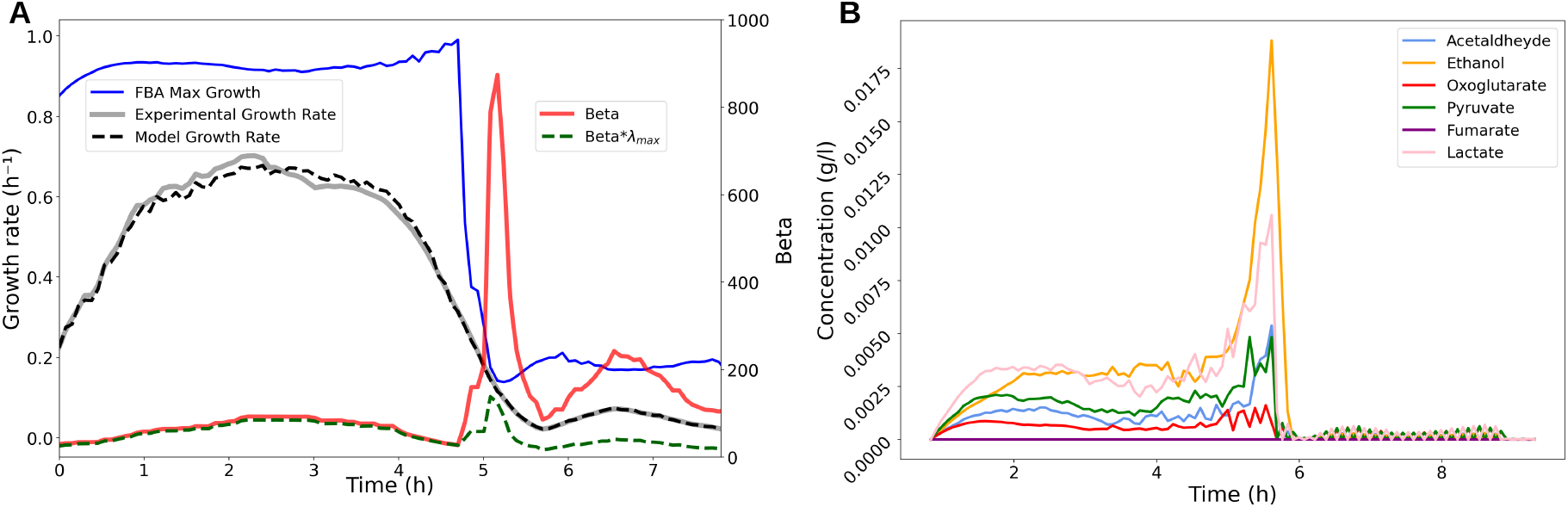
**A**. Experimental and model-predicted growth rates compared to maximal biomass production from FBA, together with the inferred time-dependent bias parameter *β*_*λ*_ in the dynMaxEnt framework. **B**. Predicted time evolution of extracellular metabolite concentrations, highlighting transient overflow-associated byproduct secretion during the metabolic transition.

Consistently, the model predicts a transient overflow metabolism phase (Fig. 8B), characterized by short-lived accumulation of fermentation byproducts such as ethanol and other organic acids during the transition. These compounds rapidly return to low concentrations once the system settles into the post-shift metabolic state, reflecting the reorganization of carbon flux distribution under the updated constraints.

## Appendix D: Single-cell metabolic flux analysis

The set of 10,000 single cell metabolic fluxes were further analyzed to identify trends in the population of fluxes. Variations in glucose and acetate exchange fluxes were present, with an inverse correlation between the two carbon imports, suggesting that the two carbon sources are largely imported under opposed metabolic conditions (Fig. 7A). Notably, acetate exchange was inversely correlated with biomass production (Fig. 7B), while glucose exchange was positively correlated with biomass production (Fig. 7C).

The distribution of metabolic states within the set of single cell metabolic fluxes was analyzed using principal component analysis (PCA) and clustering heatmap approaches. Principal component analysis highlighted a continuous distribution of cell fluxes, rather than discrete and separable modes of metabolism within the population (Fig. 7D). Principal components 1 and 2 (PC1 and PC2, respectively) were visualized in scatter plots with each point colored by major metabolic fluxes, to identify trends in major fluxes with PC1 and PC2. Acetate exchange flux showed strong correlation with PC1 (Fig. 7E). Conversely, plotting PC1 and PC2 with each point colored by glucose exchange (Fig. 7F) or biomass flux (Fig. 7G) showed strong anti-correlation with PC1.

Investigating the loadings for principal component 1 (PC1) highlighted that the switch between acetate and glucose metabolism was a key determinant for PC1. Notably, reactions with the most positive loadings for PC1 included H^+^ exchange, acetate exchange, and the reactions ACt2r and ACKr involved in acetate transport and conversion to pyruvate. In contrast, reactions involved in glycolysis and glucose fermentation, including GAPD, ENO, PDH, and glucose exchange were among the reactions with the largest negative loadings for PC1 (Fig. 7H). PC1 loadings were then visualized using the Escher core metabolism map for the e coli core model. Overlaid on the core then visualized using the Escher core metabolism map for the e coli core model. Overlaid on the core metabolism map, the trend toward enrichment of the acetate and glucose pathways was readily apparent (Fig. 7I). Thus, the PCA loadings were consistent with the finding that the switch between acetate and glucose metabolism serves as the defining axis for variance in this dataset.

**FIG. 9.**
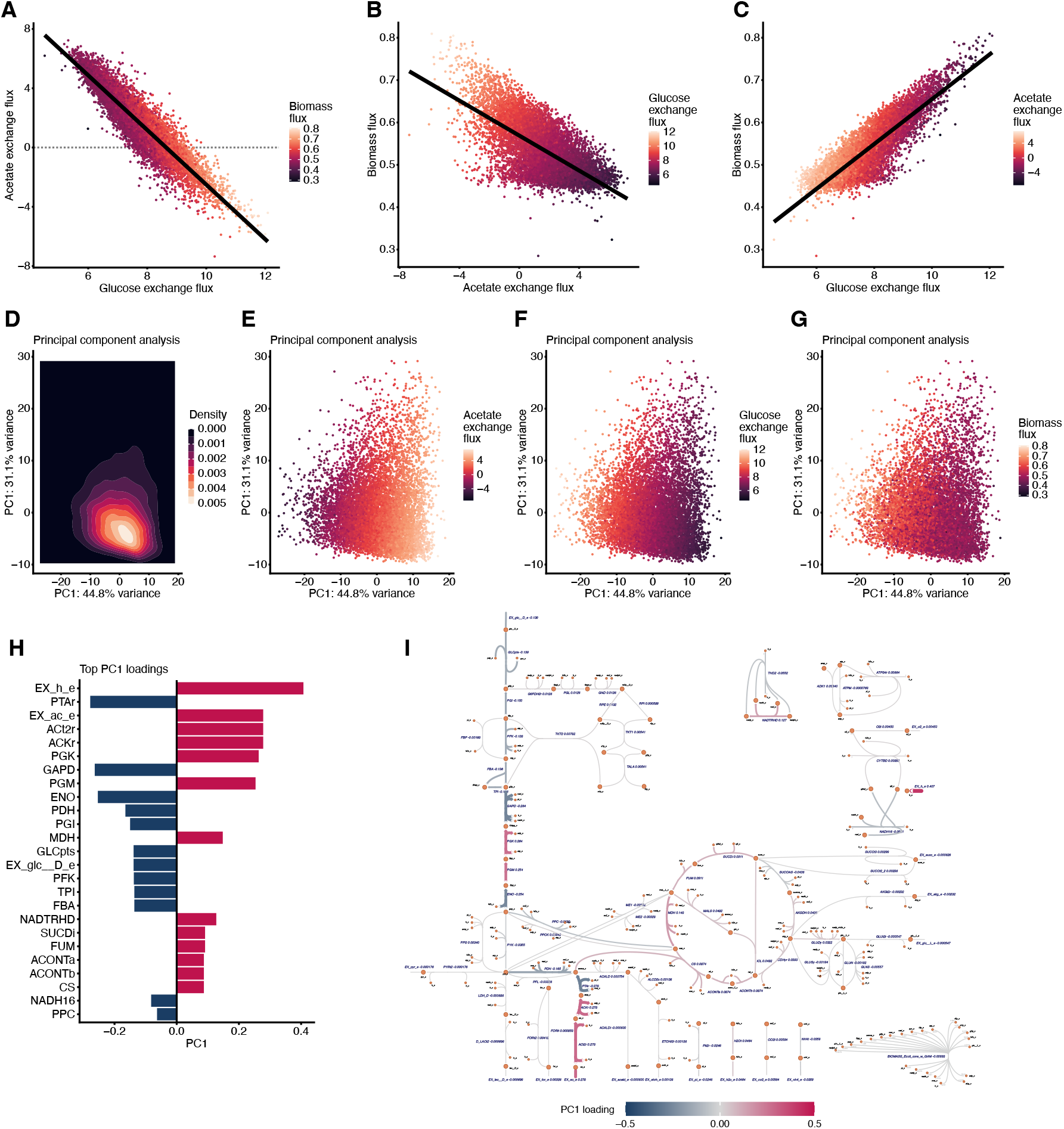
Metabolic flux analysis of single cell dataset. **A**. Scatter plot of glucose exchange flux (EX_glc_ _ D_e) and acetate exchange flux (EX ac e) in single cell data. Points are colored by biomass flux. **B**. Scatter plot of acetate exchange flux and biomass flux in single cell data. Points are colored by glucose exchange flux. **C**. Scatter plot of glucose exchange flux and biomass flux in single cell data. Points are colored by acetate exchange flux. **D**. 2D density plot for principal component analysis (PCA) values for single cell fluxes, plotted using first two principal components (PC1 and PC2). **E, F, and G**. PCA scatter plot using first two principal components, with points colored by acetate exchange flux (**E**), glucose exchange flux (**F**), or biomass flux (**G**). **H**. Top 25 reactions by loading for principal component 1. Red denotes positive value, while blue denotes negative value. **I**. Metabolic flux map of *E. coli* central metabolism, with fluxes shaded and scaled based on PC1 loadings from **H**.

### 1. Algorithmic implementation

In this subsection we briefly sketch the algorithm we use to perform the single-cell flux analysis described in the main text; an open-source implementation is available together with this work. The aim is to draw, for each of the *N*_*c*_ ∼ 10^4^ cells, a flux vector 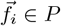 from a Boltzmann distribution

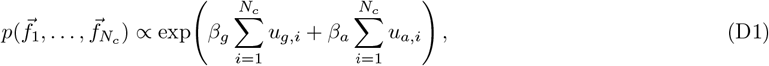

under the cell-specific constraints λ_*i*_ ∈ [λ_*i*,exp_ -*δ*_*i*_, [λ_*i*,exp_ + *δ*_*i*_], with the two Lagrange multipliers *β*_*g*_, *β*_*a*_ tuned so that the population averages 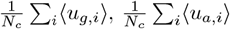 match the experimentally measured uptakes.

The implementation exploits the observation that all cells share the same underlying flux polytope P, and differ only by two scalar inequality constraints (the growth-rate window). Concretely:

1. The polytope *P* is read once from the stoichiometric matrix and the global flux bounds; an ellipsoidal rounding of *P* is precomputed and its principal axes are stored, in order to use coordinate hit-and-run on the rounded polytope, which dramatically reduces autocorrelation times [46, 47].
2. For each cell *i* = 1, …, *N*_*c*_, the global biomass bounds are tightened to [[λ_*i*,exp_ -*δ*_*i*_, [λ_*i*,exp_ + *δ*_*i*_], while all other bounds are left unchanged. A feasible interior point for cell *i* is generated by a Minover-like relaxation [75] starting from a generic initial condition.
3. Sampling proceeds in parallel over all *N*_*c*_ cells: at each Monte-Carlo sweep, an ellipsoidal hit-and-run move is proposed independently for each cell, and accepted with the Boltzmann weight 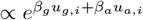 along the chosen direction (sampled exactly from the truncated exponential along the chord). To improve mixing, the multipliers are linearly annealed from 0 to their target values during a warm-up phase of order 10 sweeps.
4. The Lagrange multipliers (*β*_*g*_, *β*_*a*_) are themselves updated by a simple Boltzmann learning loop, executed manually outside the sampler: at each iteration the sampler is run to convergence, the population-averaged uptakes *ū*_*g*_(*β*), *ū*_*a*_(*β*) are estimated, and the multipliers are updated by gradient descent on the squared mismatch with the experimental targets [41, 76, 77], 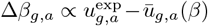 . A handful of outer iterations are typically sufficient to reach a target precision of a few percent.

After convergence, the resulting ensemble of *N*_*c*_ samples provides single-cell flux distributions consistent with the global mass-balance, the cell-specific growth-rate constraints, and the population-level uptake averages, and is the dataset used to produce Figures 3D–G. Because the cells are coupled only through the population averages defining the gradient signal in step 4, the per-sweep cost is essentially that of a single MaxEnt sampler scaled by *N*_*c*_ and is straightforwardly parallelized. The whole inference can be run on a standard workstation in a few hours for *N*_*c*_ ∼ 10^4^ and a core *E. coli* model with ∼ 100 reactions.

## Appendix E: Continuous consumer–resource model

### A. 1d continuous consumer resource model

Let us consider as a first example the case of a bacterium colony eating from a single source, glucose, in a chemostat. In this toy model we assume that:

- The growth rate *λ* and the glucose uptake *u*_G_ of a cell are proportional *λ* = *γ*_*G*_*u*_*G*_
- At optimality, the glucose uptake *U*_*G*_ depends on the glucose concentration *G* through the Michaelis-Menten Monod law 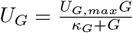 . Equivalently, we assume that the optimal growth *λ*_*0*_ is given by the law

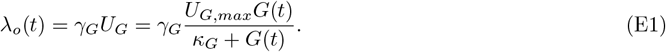

Letting ρ, *G* be the bacterium and glucose density, *p* the bacterium probability distribution, *D* the dilution rate and *G*_0_ the level of glucose in the feed, our model is the following system of two ODEs and one PDE,

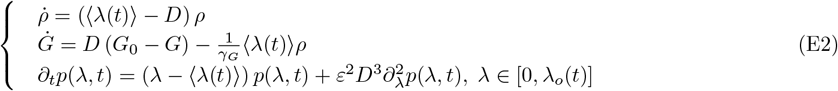

In the above system, ε > 0 is a diffusion parameter, and the average growth ⟨ λ(t) ⟩ is equal to 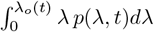. It is important to remark that in the diffusionless limit ε → 0 the probability distribution is at any moment of time a delta function centered at the value λ_*o*_, whence the system (E2) reduces to the well-known standard chemostat model

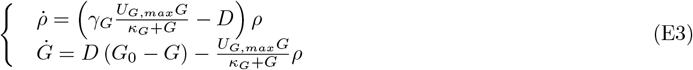

For a solution of the PDE (E2) to be admissible, the total probability must be constant in time and equal to 1. For this reason, as explained in the main text, one must impose the so-called insulating boundary conditions (12). In the 1-D case, they have a simpler form

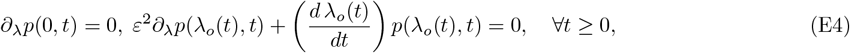

In fact, using (E4), one computes

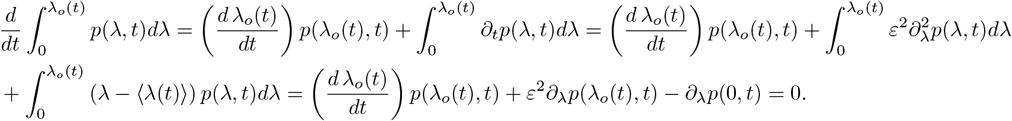

The system (E2) is called a free-boundary (or moving boundary) problem, since the domain [0, λ_*o*_(*t*)] of the solution function *p* is not known a priori, but its determination is part of the problem too. To see this at work, we look for a non-trivial fixed point of the model. This is, by definition, a **positive** bacterium and glucose densities *ρ*_*eq*_ > 0, *G*_*eq*_ > 0, and a stable distribution *p*_*eq*_ such that 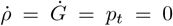 . If we are at such a point, the first equation of the system implies ⟨λ⟩ = *D*. With this information, we can solve the third equation of the system to obtain the stable distribution *p*_*eq*_(λ; ε) and its domain of definition. We change the variable to put the third equation into a standard Airy form

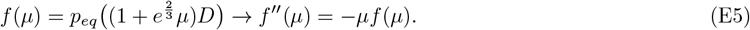

The existence of the stable distribution is equivalent to the existence of a solution to the following free boundary problem:

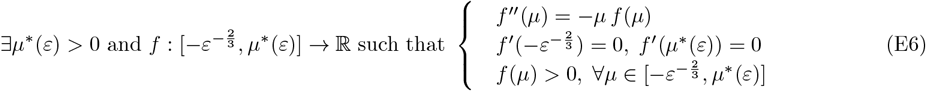

**FIG. 10.**
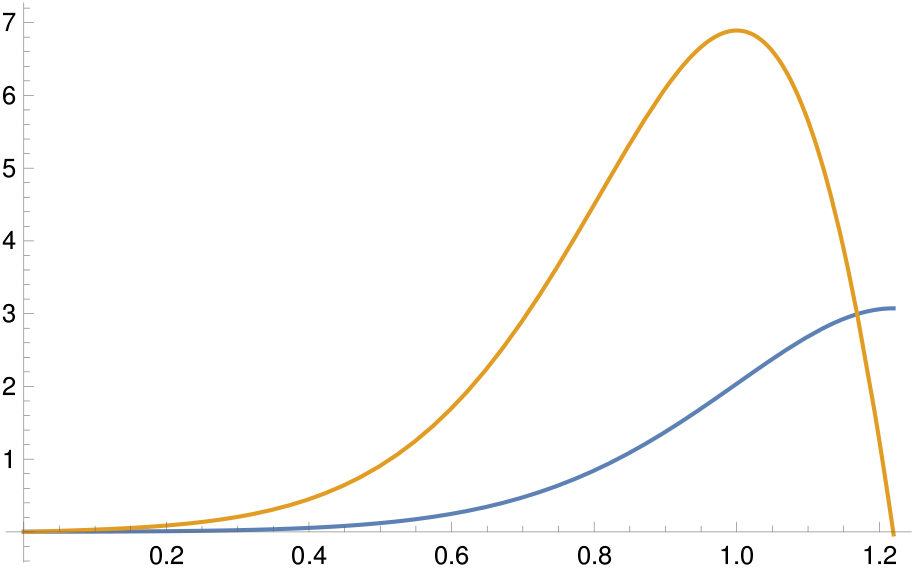
In blue the normalised solution of the free boundary problem (E6) for *ε* = 0.1, in yellow its derivative distribution has domain [0, *µ*^*\**^] with *µ*^*\**^ *≈* 0.21949. Notice that this value 0.21949 … coincides up to 5 digits with the value 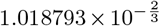 predicted by the asymptotic formula (E7).

One can show that the above problem indeed admits a unique solution for every *ε* > 0, and that the unknown *µ*^*^(*ε*) satisfies the following properties

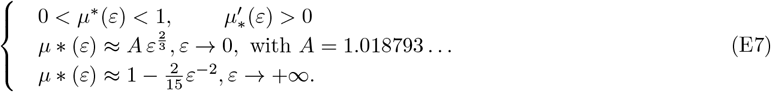

We omit the (complicated) explicit asymptotic formulas for the stable distribution, which is shown in Figure E 1 at *ε* = 0.1. Recall that the optimal growth λ_*o*_ at a given moment of time is the upper bound of the admissible λ’s at that time. In particular, at the equilibrium, the optimal growth rate coincides with the upper limit of the domain of the stable distribution, *p*_*eq*_. Using (E5), λ_*o,eq*_ is seen to be equal to 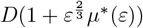. By (E1), we obtain the glucose concentration *G*_*eq*_, and, eventually, solving the second equation of the system (E2), we compute the bacteria density *ρ*_*eq*_:

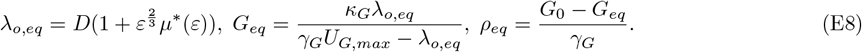

In order for these values to define a meaningful point fixed point it is necessary and sufficient that the above values of *ρ*_*eq*_ and *G*_*eq*_ are positive. Hence, we deduce an *ε*-dependent conditions on the parameters *D, G*_0_, *κ*_*G*_, *U*_*G,max*_ for the existence of the non-trivial fixed point: the non-trivial fixed point exists if and only if

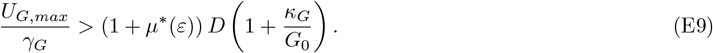

Notice that in the *ε* → 0 limit, once recovers the well-known constraint on the existence of the non-trivial fixed point for the chemostat model (E3), namely 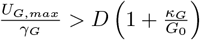 . When considering the existence of the non-trivial fixed point, the effect of the diffusion ε > 0 can be thought of as forcing a larger effective dilution rate.

### 2. Simplified *E coli* metabolic network for continuous consumer resource model

Let us now discuss a more realistic model. The state of a bacterium is determined by the glucose, acetate and oxygen fluxes *u*_*G*_, *u*_*A*_, *u*_*O*_, and we study the time evolution of the glucose, acetate, and bacteria concentration *G, A, ρ*, as well as the bacteria probability distribution *p*(*x, t*), *x* = (*u*_*G*_, *u*_*A*_, *u*_*O*_). Following the general discussion outlined above, we are led to the following free-boundary problem:

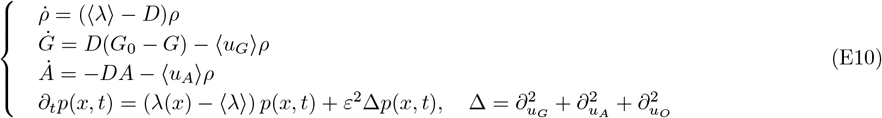

where ⟨*λ*⟩,⟨*u*_*G*_⟩,⟨*u*_*A*_⟩ are the average growth rate, glucose flux, and acetate flux. In our model the growth-rate λ is a linear function of the glucose-,acetate- and oxigen-fluxes, *u*_*G*_, *u*_*A*_, *u*_*O*_. In fact, the *carbon mass balance* implies that

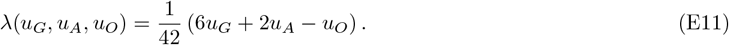

The fluxes *u*_*G*_, *u*_*A*_, *u*_*O*_, are constrained by biochemical and thermodynamic laws. In our model, we consider the following seven such restrictions

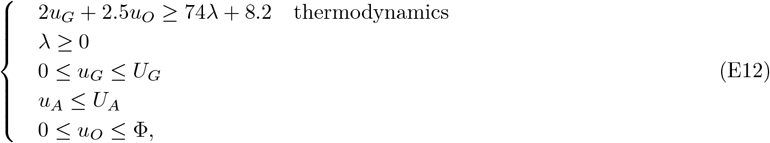

where

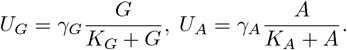

For sake of definiteness, we will later fix the constants Φ, γ_*G*_, γ_*A*_, κ_*G*_, κ_*A*_ to the experimentally realistic values

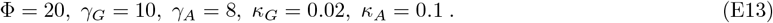

The set of constraints (E12) defines a polytope Ω(*t*) of all the fluxes which satisfy it; such a polytope depends on the parameters *U*_*G*_, *U*_*A*_, and thus on time. In order for the total probability be conserved, we must again use insulating boundary conditions. As explained in the main text, they read

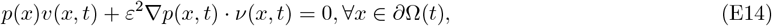

where ∂Ω(t) is the boundary of the polytope, *v* the outer normal vector, and v the outer normal velocity of the boundary.

### 3. Polytopes’ landscape

We now study how the polytope Ω depends on the values of *U*_*G*_, *U*_*A*_ as *U*_*G*_, *U*_*A*_ vary in the rectangle 0 ≤ *U*_*G*_ ≤ γ_*G*_, *0* ≤ *U*_*A*_ ≤ γ_*A*_. For later convenience, we collect all above constraints in the 7 × 4 (augmented) matrix M

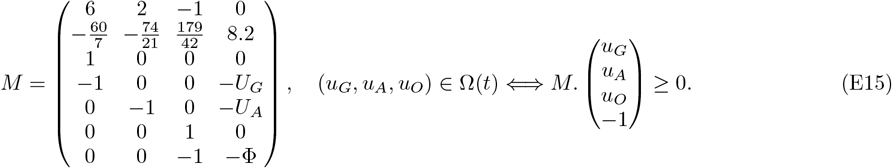

In the above formula “≥” means that all components of the vector are non-negative.

A polytope is uniquely determined by its vertices. These are the points in ℝ^3^ such that all inequalities are satisfied and at least three inequalities are saturated. We say that a vertex is simple if exactly three inequalities are saturated at it. If (*U*_*G*_, *U*_*A*_) is a generic point, there exists a neighbourhood of it such that every vertex of the polytope is simple; in such a neighbourhood all vertices vary smoothly, whence the topology of the polytope is constant. We call chamber a connected region of the *U*_*G*_, *U*_*A*_ rectangle such that all vertices of the polytope are simple. For all we said the topology of the polytope is constant inside a chamber. In order to study the chambers, it is convenient to compute their boundaries, which we call walls. There are the lines in the *U*_*G*_, *U*_*A*_ rectangle such that at least a vertex is not simple. Since a vertex is not simple if and only if at least four inequalities are saturated, a point in a wall must necessarily be a value of (*U*_*G*_, *U*_*A*_) such that a 4 × 4 minor of the matrix *M* vanishes. Since these minors depend linearly on *U*_*G*_ and *U*_*A*_, the walls are segments of straight lines. In fact, a computation (that we omit) shows that the walls are the segments of the following 7 lines which lie the *U*_*G*_, *U*_*A*_ rectangle :

**FIG. 11.**
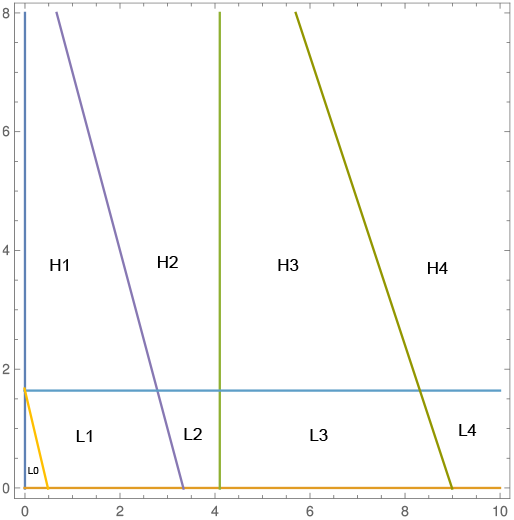
Chambers and Walls of the *U*_*G*_, *U*_*A*_ rectangle.

**FIG. 12.**
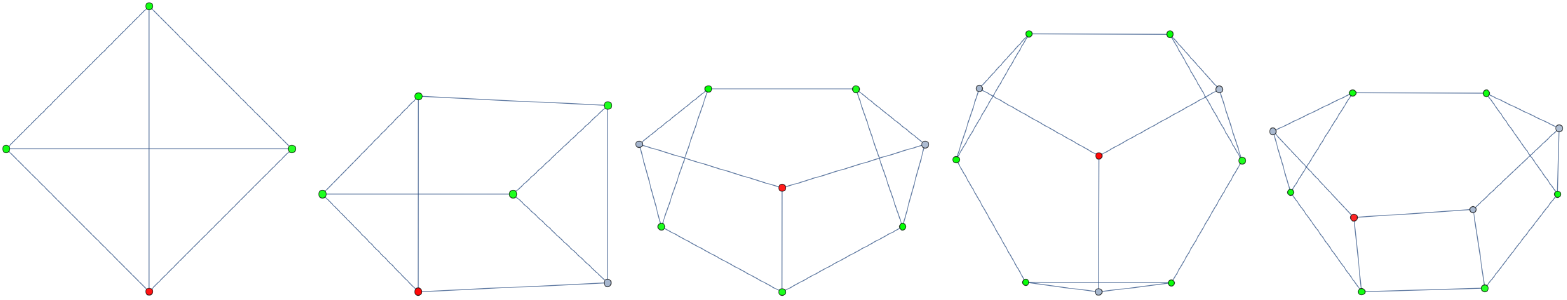
The topological types of the graphs associated with the cells, ordered with respect to number of vertices. The first type corresponds to chamber *L*_1_, the second type corresponds to chambers *L*_2_, *H*_1_, the third type corresponds to chambers *H*_2_, *L*_3_, *L*_4_, the fourth type corresponds to chamber *H*_3_, and the fifth type to chamber *H*_4_. In red, the vertex optimizing the growth *λ*. In green, the vertices at which the growth *λ* is minimal, namely *λ* = 0. Notice that the growth *λ* is zero on the whole face spanned by these vertices (since it is a linear function).

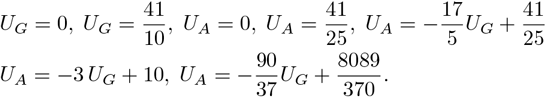

These lines partition the rectangle into 9 chambers *H*_1_, *H*_2_, *H*_3_, *H*_4_ and *L*_0_, *L*_1_, *L*_2_, *L*_3_, *L*_4_ according to the figure E 3 (*H* stands for high acetate density and *L* for low acetate density). *L*_0_ is the unfeasible region, that is the connected component of the *U*_*G*_, *U*_*A*_ rectangle such that the polytope is empty - no set of fluxes satisfies the inequalities.

The topological type of the polytopes belonging to each chamber are shown in Figure E 3 below. Notice that a few chambers share the same topological type. This means that when crossing a wall some vertices coalesce and others pop-up, the new polytopes that arise may be topologically equivalent (however, if one decorates the vertices with the triple of inequalities that are saturated, then the decorated polytopes are different in different chambers).

Since the growth λ is a linear function, then its optimal value is attained at a vertex of the polytope. A computation (that we omit) shows that inside each chamber a unique vertex is optimal and such a vertex is constant in the chamber. Which is the optimal vertex? As we remarked above, a vertex is defined by the saturation of three inequalities. Our computations show that at the optimal vertex two of the three inequalities saturated at the optimal vertex are the same in every chamber, namely the glucose flux is maximal *u*_*G*_ = *U*_*G*_ and the thermodynamical bound 74λ = 2*u*_*G*_ + 2.5*u*_*O*_ − 8.2 is saturated. Moreover, we observe that the third saturated inequality at the optimal vertex is either the acetate flux, i.e. *u*_*A*_ = *U*_*A*_, or the oxygen flux, i.e. *u*_0_ = Φ. In other words, varying *U*_*G*_ and *U*_*A*_ an acetate switch occurs: in the polytope landscape, a line separate the region where the optimal growth is attained when the oxygen flux is maximal from the region where the optimal growth is attained when the acetate flux is maximal. Such a line is given by formula 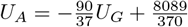 .

**FIG. 13.**
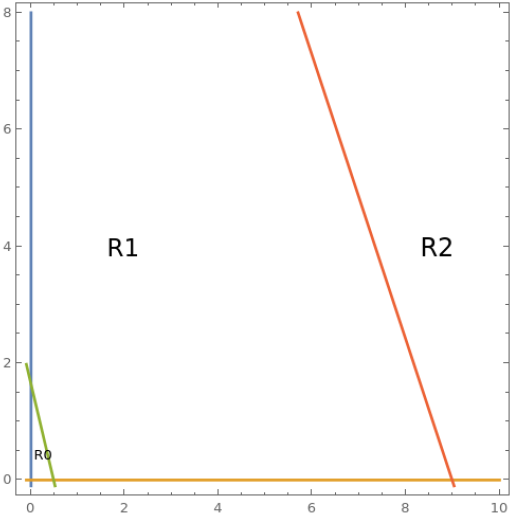
The regions *R*_0_, *R*_1_, *R*_2_ in the *U*_*G*_, *U*_*A*_ rectangle. Region *R*_0_ is the unfeasible region; in region *R*_1_, the optimal growth implies that the acetate flux is maximal; in region *R*_2_, the optimal growth implies that the oxygen flux is maximal while the acetate flux may be even negative. Region *R*_0_ and *R*_1_ are separated by a segment of the line 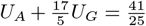, while region *R*_1_and *R*_2_ are separated by a segment of the line 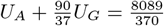.

More precisely. we observe that, with respect to optimal growth, the *U*_*G*_, *U*_*A*_ rectangle divides in only three regions, see Figure E 3:

- The unfeasible region *R*_0_ = *L*_0_.
- The region R_1_ = H_1_∪ H_2_ ∪H_3_∪ L_1_∪ L_2_∪ L_3_ in which the acetate flux is saturated at optimal vertex, that is *u*_*A*_ = *U*_*A*_ ; in this region, the optimal growth is

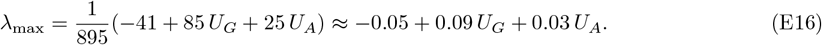

Notice that, when *U*_*A*_ = 0 and U_*G*_ is not small, the above law is consistent with the simple 1-D chemostat model, in which λ_max_ ≈ 0.1 *U*_*G*_.
- The region *R*_2_ = *H*_4_ ∪*L*_4_ in which the oxygen flux is saturated at the optimal vertex, that is *u*_*O*_ = Φ; in this region, the optimal growth is

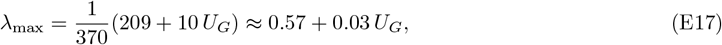

and the acetate flux is

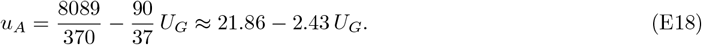

Taking into consideration the acetate flux, we divide the region R_2_ into two distinct regions 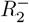, where the acetate flux is negative, and 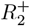 where the acetate flux is positive. By above formula, the region 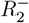 is simply the rectangle 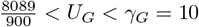 and 0 ≤ *U*_*A*_ ≤ γ_*A*_ = 8.

While the optimal value of the growth is attained at a single vertex, on the contrary its minimal value, namely λ = 0, is attained on at least three vertices which span a face of the polytope on which the growth is zero; see Figure E 3.

### 4. Diffusionless limit

A thorough analysis of the system (E10) is well beyond the scope of this paper. We consider here merely its diffusionless limit ε → 0, when the probability distribution is at any moment of time a delta function centered at the optimal value of the growth rate. In this limit the PDE drops out of the system, and we are left with the system of ODEs

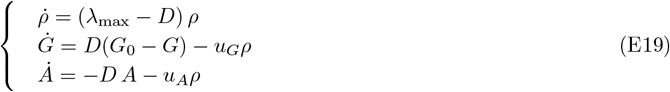

The quantities λ_max_, *u*_*G*_, *u*_*A*_ are functions of *G, A* via *U*_*G*_, *U*_*A*_ and their values is defined by their values at the optimal vertex. Therefore, after the discussion above, we have

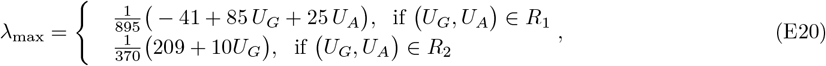

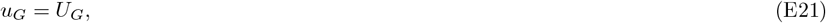

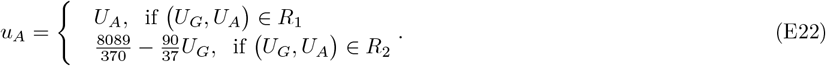

We conclude our analysis with the study the fixed points of the diffusionless system (E19). Fixed points are, by definition, those values of *G*_*eq*_, *ρ*_*eq*_, *A*_*eq*_ such 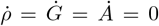. We say that a fixed point is non-trivial when *ρ*_*eq*_ > 0.

*Non-trivial fixed point with null acetate concentration* In this case *ρ*_*eq*_ > 0 but *A*_*eq*_ = 0. This fixed point lies in the region R_1_. It exists for the following values of the parameters *D, G*_0_:

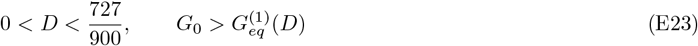

With

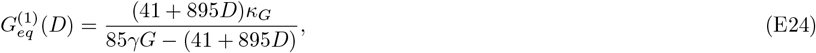

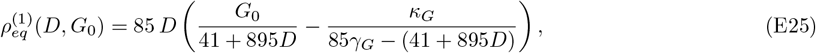

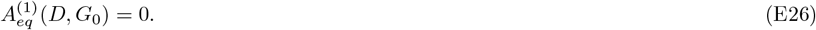

The fixed point is stable whenever it exists. See Figure E 4.

*Non-trivial fixed point with positive acetate concentration (negative acetate flux)* In this case *ρ*_*eq*_ > 0 as well as *A*_*eq*_ > 0. This fixed point lies in the region 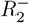. It exists for the following values of the parameters *D, G*_0_:

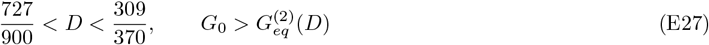

With

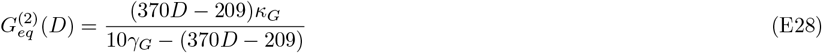

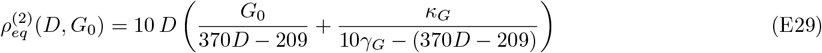

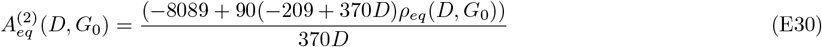

The fixed point is stable whenever it exists.

**FIG. 14.**
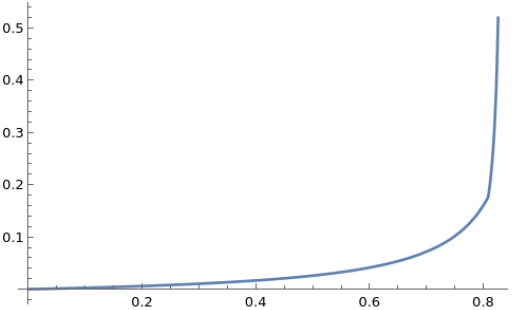
In the plot, the curves *G*_*eq*_ (*D*) depending on the flux *D* for the *D*-range 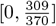 such that a non-trivial fixed point exists. The admissible *D* range is divided in two segments, 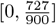 where the non-trivial fixed point has null acetate concentration, and 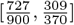 where the non-trivial fixed point has non-trivial acetate concentration (and negative acetate flux). In the first segment *G*_*eq*_ (*D*) is given by the function 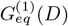, while in the second segment *G*_*eq*_ (*D*) is given by the function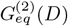.If *D* is outside the range 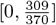 or *G*_0_ *< G*_*eq*_ (*D*) then the trivial fixed point *ρ* = *A* = 0, *G* = *G*_0_ is stable.

### 5. Moran-like particle scheme for the 3D model

The full 3D model (E10), augmented by the regulatory advection term introduced in the main text and by the time-dependent polytope Ω(t), is a moving-boundary nonlinear PDE for which standard finite-difference schemes are awkward to set up. We therefore solve it with a heuristic particle method that is straightforward to implement, scales easily to higher dimensions, and reproduces with good accuracy the analytical predictions in the cases where they are available.

The state of the population is represented by *N*_*p*_ ∼ 10^4^ tracer particles 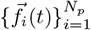, each carrying a 3D flux vector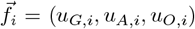 that lies inside the current polytope Ω(*t*). At each time step Δ*t* the algorithm performs three elementary processes followed by a constraint update. The implementation we used corresponds to the open-source code released with this paper.

1. **Biased diffusion (advection + Brownian motion)**. For each particle a candidate displacement of size 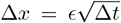 is proposed along one of the ± flux axes (chosen uniformly at random). The proposed move is rejected outright if it leaves Ω*(t*), and otherwise accepted with a Metropolis weight min( 1, *e*^ΔΦ^), where ΔΦ = *β* Δλ + *β*_*g*_ Δ*u*_*G*_ + *β*_*a*_ Δ*u*_*A*_. Setting *β* > 0 implements the regulatory advection toward higher λ discussed in the main text [58], while *β*_*g*_, *β*_*a*_ are reserved for the maximum-entropy bias matching exchange averages whenever needed.
2. **Replication–death (Moran step)**. Each particle attempts to replicate with probability λ_*i*_ Δ*t*. To keep the total number N_*p*_ of tracers strictly constant—and thus enforce the chemostat-level normalization of *p*(*x, t*) *out of equilibrium* too—each successful replication is paired with the elimination of a uniformly chosen particle, which is overwritten by the offspring’s state. This is exactly a generalized Moran step [78, 79] with selection driven by the local growth rate; in the continuum limit it reproduces the replicator term (λ(*x, E*) λ ) *p*(*x, t*) in (11).
3. **Environmental update**. Population-averaged uptake fluxes ⟨*u*_*G*_⟩,⟨*u*_*A*_⟩ and growth rate ⟨λ⟩ are computed from the current ensemble, and the chemostat ODEs for *G, A, ρ* are advanced by an explicit Euler step using Δ*t*.
4. **Constraint update with fixed-front rescaling**. The new nutrient concentrations modify the upper bounds *U*_*G*_(*G*), *U*_*A*_(*A*) and thus reshape the polytope Ω(*t* + Δ*t*). To keep particles inside the moving domain we adopt a heuristic in the spirit of fixed-front methods [59]: each flux coordinate is rescaled by the ratio of new and old bounds, 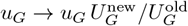, with analogous rules for *u*_A_, *u*_O_ that preserve the topology of Ω as far as possible. Particles that would still violate the new constraints after this contraction/expansion are reinjected uniformly into the interior of Ω(t + Δt) via rejection sampling.

This scheme is heuristic: in particular, the rescaling step in (4) is not derived from the underlying PDE but is justified *a posteriori* by its accuracy. As a benchmark, in the 1D version of the problem (Appendix E 1) we have implemented an independent Crank–Nicolson finite-difference solver with explicit moving-boundary tracking. The two methods give numerical results in close agreement and are both consistent with the analytical asymptotics derived in Appendix E 1 (see Fig 14 below), which provides confidence in the use of the particle scheme also in the higher-dimensional setting where direct PDE methods become impractical.

**FIG. 15.**
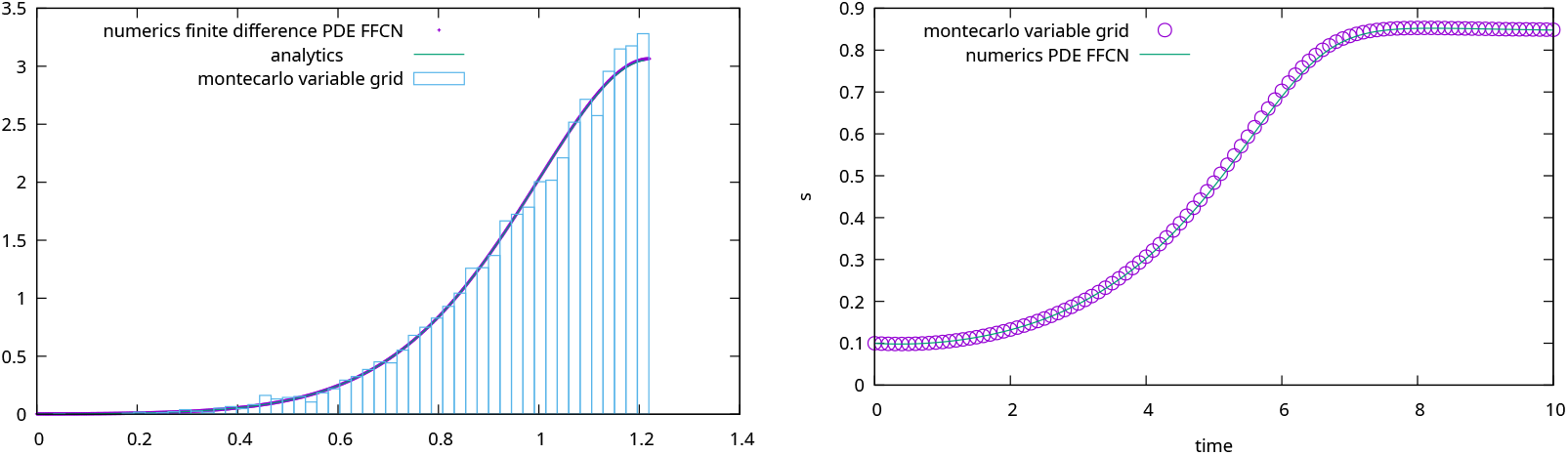
1d Continuous consumer resource model. Left: steady state distribution of phenotypes, comparison among solutions obtained with the analytical approximation, the finite difference numerical algorithm and the Montecarlo method. Right: density as a function of time, comparison between the finite difference numerical algorithm and the montecarlo method. Calculations are performed for the case *ϵ* = 0.1.

